# Physiological and transcriptomic characterization of cold acclimation in endodormant grapevine under different temperature regimes

**DOI:** 10.1101/2023.10.21.563432

**Authors:** Hongrui Wang, Al P. Kovaleski, Jason P. Londo

## Abstract

It is essential for the survival of grapevines in cool climate viticultural regions that vines properly acclimate in the late fall and early winter and develop freezing tolerance. Climate change-associated abnormities in temperature during the dormant season, including oscillations between extreme cold and prolonged warmth, impacts cold acclimation and threatens the sustainability of the grape and wine industry. We conducted two experiments in controlled environment to investigate the impacts of different temperature regimes on cold acclimation ability in endodormant grapevine buds through a combination of freezing tolerance based physiological and RNA-seq based transcriptomic monitoring. Results show that the freezing tolerance of buds was not altered from field levels when exposed to stable temperatures ranging from 2 °C to 22 °C but was enhanced when exposed to temperature cycling (7±5 °C). We also characterized the transcriptomic response of endodormant buds to high and low temperatures and the potential genetic control for the maintenance of endodormancy. Several pathways that were previously reported to be responsive or functional during cold acclimation, such as the *ICE-CBF-COR* cascade, were not observed to play a role in the enhancement of freezing tolerance or the sensing of different temperatures, indicating our current understanding of the genetic control of cold acclimation remains a challenge when generalizing across plant species and phenological stages.

## 1 Introduction

Climate change, a phenomenon attributed to direct or indirect human activities, is characterized by long-term changes of climate status and unseasonable weather events ^1–3^. As a consequence, the steady movement of optimum planting zones towards polar regions and more frequent weather extremes pose significant challenges to food production, particularly to the production of perennial crops such as grapevine, due to high reestablishment cost and slow breeding processes ^4–6^. Other than an expected migration of vineyards towards cooler regions and a shift to more adaptive cultivars in the future, the grape industry is currently experiencing major crop losses due to climate change-associated weather extremes such as extreme drought, precipitation, heat and cold ^7–12^. Among abiotic stresses, cold-related damage in winter and spring is one of the leading constraining factors for the expansion of viticulture into cool and cold climates. Therefore, understanding adaptations to cold are key to the development of management methods for cold damage for the sustainability of grape and wine production.

To overcome temperature adversity in winter, perennials like grapevines develop freezing tolerance through cold acclimation ^13^. As a three-phase process, cold acclimation is triggered by short-day photoperiod in late summer and step-wise enhanced by low above-freezing and sub-freezing temperatures ^14,15^. In general, the enhancement of freezing tolerance during this process is a largely unresolved consequence of sophisticated combination of genetic control, metabolism amelioration and physiological adaption ^16,17^. Genetically, key regulatory pathways or cascades, such as the abscisic acid (ABA) signaling pathway, the ethylene signaling pathway, the jasmonate signaling pathway and inducers of *CBF* Expression – C-Repeat Binding Factor/*DRE* Binding Factor – Cold Regulated Genes (*ICE-CBF/DREB1-COR*) cascade (*ICE-CBF-COR*), are activated to initiate the expression of functional proteins and metabolites ^18–22^. Metabolically, functional metabolites and proteins such as soluble sugars, proline, reactive oxygen species-detoxification proteins, ice binding proteins, flavanols and anthocyanins, accumulate in cells of overwintering buds ^17,23–25^. Physiologically, adaptive mechanisms such as cell desiccation, modification of cellular lipid composition, glass formation and supercooling, are thought to develop to either avoid or tolerate intracellular ice formation, which in turn enhances freezing tolerance ^26–28^.

In parallel to cold acclimation, dormancy status also shifts from paradormancy during the growing season (inhibition of growth caused by apical dominance) to endodormancy (inhibition of growth caused by internal molecular constraint) ^29^. Endodormant buds in woody perennials remain unbroken even under growth-permissive conditions, until transitioning to an ecodormant state (inhibition of growth due to environmental conditions), which occurs after being exposed to sufficient chilling length ^30,31^. The response to chilling accumulation varies among *Vitis* species ^32^. Although many studies have explored aspects of endodormancy, the mechanism of growth inhibition is not understood. *DORMANCY-ASSOCIATED MADS box genes* (*DAMs*) were noted for their potential functionality during endodormancy. A locus related to the deletion of tandemly repeated *DAMs* resulted in ever-growing peach without dormancy ^33^. In *Prunus persica*, the expression of *DAM1-4* peak at bud set and might function on the initiation of endodormancy, while the expression of *DAM5* and *DAM6* peak at the start of endodormancy and steadily decrease as being exposed to prolonged chilling, suggesting their potential function in maintaining endodormancy ^34–36^. Functional studies suggest that the conservation of amphiphilic repression motif in tandemly arrayed *DAMs* might lead to direct transcription repression, and *DAMs* might also interact with other dormancy-promoting hormones or proteins ^37–39^. Recent studies reported that the regulation of *DAM* expression is achieved by chromatin covalent modifications, DNA methylation or miRNA, suggesting the involvement of epigenetic control in *DAM*-mediated dormancy ^40–42^. In grapevine, homologs of flowering time control genes in *A. thaliana* and other model species were identified, and some these genes, including some *MADS box* genes, were reported to highly express during dormancy and lowly express during budbreak and flowering ^43^. As climate change is predicted to lead to, on average, warmer winters with more frequent extreme warming or freezing events, grapevine survival in dormant season and proper growth after dormant season would be more challenging regarding the synergy of dormancy maintenance, chilling requirement fulfillment and sufficient freezing tolerance ^3,11,44,45^. Thorough understanding of grapevine physiology and genetic control during cold acclimation and endodormancy along with the alternation of these mechanisms under different temperature regimes is urgently needed.

In this study, we conducted two experiments to investigate grapevine cold acclimation in endodormant buds under different temperature regimes. The first experiment involved the examination of grapevine cold acclimation in dormant cuttings exposed to five stable temperatures ranging from 2 °C to 22 °C. The second experiment assessed the impact of diurnal temperature cycling through comparing grapevine cold acclimation under stable temperature environment to a cycling temperature environment. In both time-course experiments, bud freezing tolerance was measured using differential thermal analysis (DTA) in parallel with RNA-seq based transcriptome characterization to connect phenotypic change (freezing tolerance) with potential underlying genetic control. This work targets the gap of knowledge regarding grapevine cold acclimation and endodormancy and offers insight for developing novel viticultural management methods to sustain grape production in adaptation to climate change.

## 2 Materials and Methods

### 2.1 Plant material

Dormant cuttings collected from field grown *Vitis vinifera* ‘Cabernet Sauvignon’ cultivated at ‘Ravine’s Wine Cellars’ in Geneva NY (42.845° N, 77.004° W) were used for experimentation. ‘Cabernet Sauvignon’ was grafted on 3309C rootstocks and subjected to standard vineyard management. Canes were collected on 26 October 2016 (chilling unit = 398, Utah model) when grapevines were at Eichhorn Lorenz Stage 1 (EL1 – winter bud) ^46^. At this timepoint, chilling requirement is not fulfilled for ‘Cabernet Sauvignon’ ^32^, and the buds were endodormant. After field collection, canes were chopped into single-node cuttings, cuttings were randomized, and placed in cups with cut ends in water. Cuttings were stored in a walk-in cold room at 7 °C for one day until being used for experimentation. During the experiments, water was added to the sample cups periodically to maintain humidity.

### 2.2 Experiment 1: cold acclimation under different stable temperatures

Single-node cuttings were divided into five groups, and each group was subjected to a temperature treatment in growth chambers: 22 °C, 11 °C, 7 °C, 4 °C and 2 °C. These treatments were set at stable temperatures without any diurnal fluctuation and no light exposure. Sample collection for freezing tolerance assessments and RNA-seq was conducted from growth chambers at eight sample times: at 7 d, 13 d, 20 d, 26 d, 34 d, 41 d, 55 d and 69 d of growth chamber temperature exposure. Samples in the 22 °C treatment were not collected at the 69 d post-collection due to budbreak occurring in the growth chamber.

### 2.3 Experiment 2: cold acclimation under temperature cycling

Experiment 2 was conducted in parallel with Experiment 1 but was temporally separated into two phases: phase I and phase II. The schematic presentation of the experiment is shown in Figure S1. During phase I, temperature exposure of the cuttings was held stable at 11 °C, 7 °C, or 2 °C in different growth chambers starting at 0 days after field collection and continuing to 26 days after collection. At 26 d after collection, the cuttings were then moved from each of the 11 °C, 7 °C, and 2 °C stable treatments and placed into a shared growth chamber where the mean temperature was maintained at 7 °C with a diurnal cycling of ±5 °C (max = 12 °C, min = 2 °C). Cuttings were held at each temperature for 6 h, creating a 10 °C oscillation analogous of daily temperature fluctuations (6 h at 2 °C, 6 h at 7 °C, 6 h at 12 °C, 6 h at 7 °C, repeat). The treatment is abbreviated as a ‘7±5 °C’ treatment. Cuttings were maintained in the cycling chamber until 69 d after field collection. Sample collection for freezing tolerance assessments and RNA-seq were conducted in parallel with Experiment 1.

### 2.4 Freezing tolerance measurement

Differential thermal analysis (DTA) was used to determine grapevine bud freezing tolerance following standard protocols ^47,48^. During each collection, buds were excised along with surrounding tissue from single bud cuttings and loaded into thermoelectric modules in a programmable freezer to be subjected to a freezing rate of 4 °C/h from 0 °C to −50 °C. The release of heat during tissue freezing, a phenomenon known as the low temperature exotherm (LTE) was recorded via a Keithley 2700 data logger (Tektronix, Beaverton, OR, USA). The temperature at which LTE occurs corresponds to the freezing tolerance of the grapevine bud ^48,49^. Freezing tolerance was determined for each treatment and collection timepoint by recording the freeze temperature of five biological replications.

### 2.5 RNA-seq library preparation and data processing

At each collection time point, bud samples were excised from single-bud cuttings and flash-frozen in liquid nitrogen. Three biological replicates, each consisting of five pooled buds, were collected per treatment. Frozen bud samples were stored at −80 °C until extraction. The Spectrum^TM^ Plant Total RNA Kit (Sigma Aldrich, St Louis, MO, USA) was used to extract total RNA from ground bud tissues. Cornell University Institute of Genomic Diversity (Ithaca, NY, USA) provided technical support for library construction using Lexogen QuantSeq 3’mRNA-Seq Prep Kit (Lexogen, Greenland, NH, USA) following manufacturer protocol. Sequencing of libraries were conducted at Cornell University Institute of Biotechnology (Ithaca, NY, USA) using NextSeq500 (Illumina, Inc., San Diego, CA, USA) with 95 samples per lane. The read length was 85 bp, and sequencing was replicated three times in each library for technical validity.

As the first step of RNA-seq data processing, FastQC ^50^ was applied to each library for quality control. BBDuk ^51^ was employed to remove poly-A and adaptors following the default standard pipeline. STAR ^52^ was used for the alignment of trimmed sequences against the *Vitis. vinifera* 12X.v2 genome and VCost.v3 annotation ^53^. Transcript quantification at gene level was accomplished via ‘-quantMode GeneCounts’. In each experiment, all gene counts (as row) in all samples (as column) were merged as a gene count matrix. Low count genes (total gene count < total sample count) were excluded, and the remaining matrix was analyzed and normalized in DESeq2 ^54^. In Experiment 1, the full model of DESeq2 contained days in treatments and treatment temperatures as continuous variables. In Experiment 2, the full model of DESeq2 contained the interaction of phase I treatment and phase II treatment as a discrete variable. Variance stabilization transformation (VST) and gene expression normalization were also conducted in DESeq2 for downstream gene co-expression network analysis and gene expression visualization, respectively.

### 2.6 Outlier filtering with PCA and gene co-expression network analysis with WGCNA

In each experiment, VST counts of all genes were analyzed using principal component analysis (PCA) to detect outliers. The samples showing apparent deviation from the main sample cluster in the first three principal components (PCs) were identified as outliers and excluded from gene count matrix for downstream analysis. DESeq2 analysis was conducted again with the filtered gene count matrix using the same models, and resulting VST gene counts were used as input for weighed gene co-expression network analysis (WGCNA) ^55^. Co-expression network was constructed using remaining samples and genes with ‘blockwiseModules’ (power = 12, networkType = “signed”, TOMType = “signed”, minModuleSize = 50, reassignThreshold = 0, mergeCutHeight = 0.25).

### 2.7 Identification of target co-expression modules and target DEGs

After the identification of co-expression modules through WGCNA, gene members in module ‘grey’ represented “noise” and were excluded for any downstream analysis. In each remaining module, module eigengenes (MEs), were tested for their correlation to each variable in the experiment with the Pearson method ^55^. MEs of samples were also visualized across the experiment. Target co-expression modules were determined based on the p-values of MEs-variable correlation and the identification of reasonable pattern in ME visualization per interest of the experiment (e.g., temperature effect in Experiment 1 or 7±5 °C effect in Experiment 2). The genes in target modules were further subjected to correlation analysis (e.g., correlation of gene expression with temperature in Experiment 1) or contrasting (stable vs. 7±5 °C in Experiment 2) in DESeq2. The genes with FDR < 0.05 in correlation analysis or FDR < 0.05 and log2 fold change (LFC) > 2 in DESeq2 were identified as target differentially expressed genes (DEGs) and were subjected to downstream analysis.

### 2.8 Pathway enrichment analysis with GSEA

Pathway enrichment analysis of target DEGs were conducted using gene set enrichment analysis (GSEA) ^56^. To obtain better functional annotation, the identification of target genes was derived from their CRIBI V1 annotation (http://genomes.cribi.unipd.it/grape), and gene models without V1 annotation were excluded for GSEA. Remaining genes were pre-ranked using the FDR from correlation analysis or contrasting in a decreasing order. The pre-ranked gene list was transformed to a .rnk file as a part of input for GSEA. Pathway information is obtained from VitisNet database ^57^, and all pathways were combined into a gene set file (.gmt) as another part of input for GSEA. GSEA was conducted with ‘Run GSEAPreranked’ in weighted mode with 1000 permutation. Normalization mode was set as ‘meandiv’, and no max size or min size was chosen for any gene set.

### 2.9 Manual examination of top DEGs and genes of interest

In addition to pathways enrichment analysis, we manually investigated the top 200 DEGs (ranked based on LFC) that showed significance in the specific statistical analysis in each experiment, depending on the result of differential expression analysis. In Experiment 1, the top 200 DEGs that showed significant positive, or negative correlations with temperatures were examined. In Experiment 2, the top 200 DEGs that showed significance in the contrast of stable temperature vs. 7±5 °C were examined. During these manual examinations, when groups of genes were observed in the top 200 DEG list and were also repeatedly associated with known pathways the DEGs were manually analyzed and visualized for their response to the specific temperature treatments during the experiment. Some genes of interest, including the genes encoding for ICE or CBF/DREB were also subjected to manual examination.

## 3 Results

### 3.1 RNA-seq data statistics

In total, 156 libraries were sequenced in the two experiments of this study. All libraries passed the FastQC per base quality test, indicating good quality of raw reads. Mean read length per sequence of raw reads was 85 bp. After trimming to remove adaptor barcodes, the mean read length was 79 bp. The read per library averaged 4.2 million and the uniquely mapped rate per library averaged 78.9%. One library, a replicate of phase I 2 °C × phase II 7±5 °C in Experiment 2, showed significantly lower uniquely mapped rate at 10.59% and was identified as an outlier in the sample PCA analysis and removed.

In Experiment 1, the gene count matrix composed 120 samples and after low count filtering, 16,346 of the total 42,413 genes (39%) in the VCost.v3 annotation were detected. These genes were analyzed by DESeq2 to generate VST counts. PCA of VST counts identified six outliers (Figure S2) in Experiment 1, and these outliers were excluded. The remaining genes and samples were subjected to downstream analysis.

In Experiment 2, the gene count matrix composed 111 samples and after low count filtering, 16,028 genes (38%) were detected, and these genes were analyzed by DESeq2 to generate VST counts. PCA of VST counts identified six outliers, including the library that showed significantly lower uniquely mapped rate (Figure S3) in Experiment 2, and these outliers were excluded. The remaining genes and samples were subjected to downstream analysis.

### 3.2 Experiment 1: Undifferentiated phenotype but differentiated transcriptome

All buds remained dormant and unbroken during Experiment 1, with the exception of the buds treated at 22 °C at 69 d post-treatment. The freezing tolerance of buds from each collection is represented by the LTEs detected in DTA, and the means of the LTEs of each treatment × timepoint are shown in Figure 1A. The LTEs of all temperature treatments ranged between −8.0 °C to −14.5 °C during the experiment (Figure 1A). The LTEs of each treatment were analyzed by Pearson method for correlations with days in treatment (Figure 1B).

**Figure 1.**
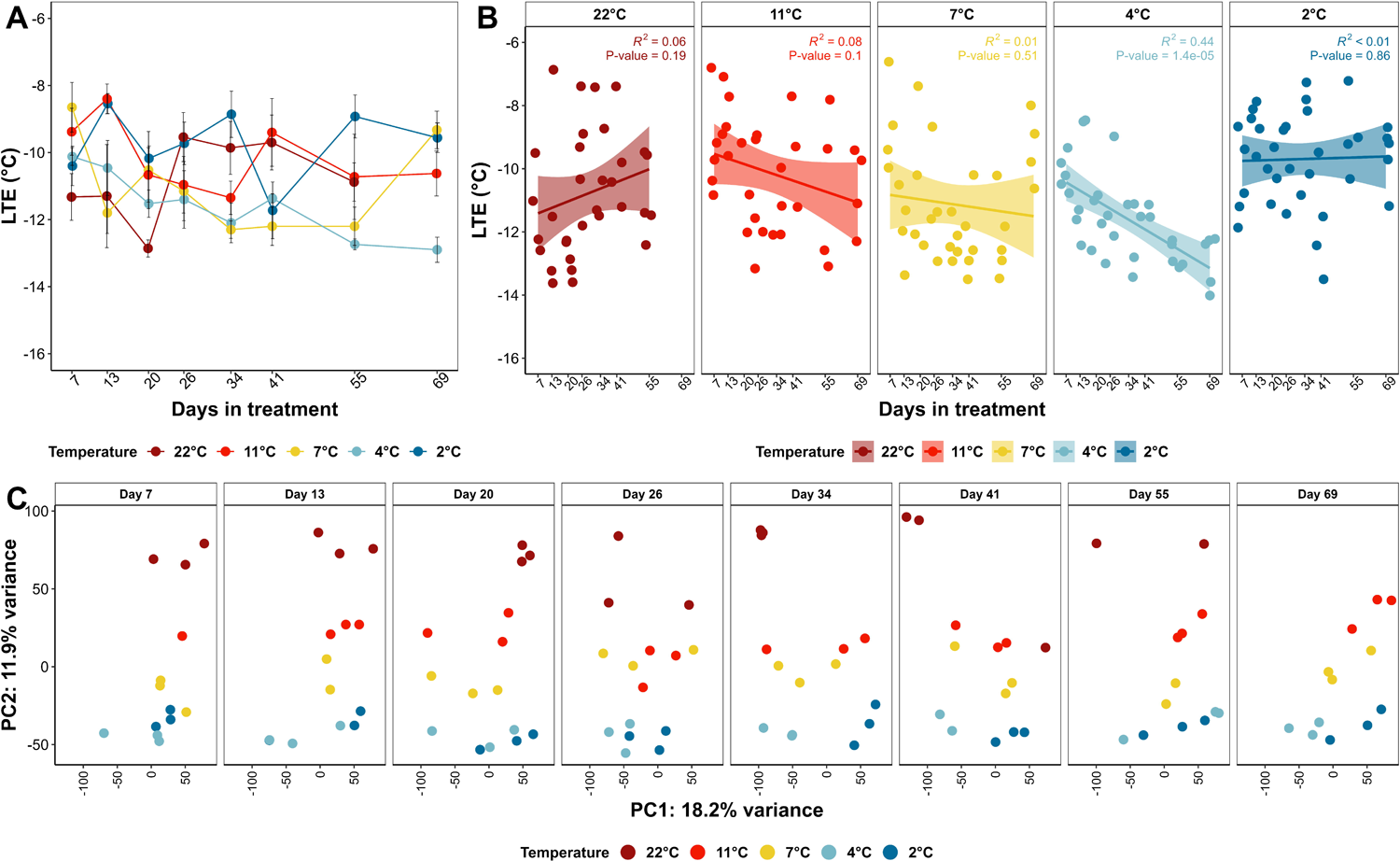
Freezing tolerance measurement and transcriptome characterization of grapevine buds in Experiment 1. A) Mean low temperature exotherm (LTE) across the experiment. Bars represent means ± SE (n = 5); B) Correlation analysis of days in treatment and LTE; C) Principal component analysis (PCA) of gene expression changes after outlier filtering.

Following the analysis and normalization in DESeq2 and outlier filtering, transcriptome characterization was conducted on VST counts of the remaining genes through PCA. The top two PCs, PC1 and PC2 explain greater than 30% of total variance, and the PC values of all the remaining samples are shown across the experiment in Figure 1C. Correlation analysis of PC value vs. temperature indicates that PC1 of the samples do not correlate with either treatment temperature or days in treatment (Figure S4A and B), whereas PC2 of the samples significantly correlate with treatment temperatures (Figure S4C).

### 3.3 Experiment 1: DEGs and enriched pathways under different temperatures

After low count filtering, VST counts of detected genes (16,346 genes) in Experiment 1 were examined with WGCNA to construct gene co-expression network and examine gene co-expression modules. In total, 24 gene co-expression modules (excluding module ‘grey’) with number of genes ranging from 63 to 2,950 were identified. The MEs were tested for correlation with days in treatment and temperature and were visualized across the experiment (Figure S5). Among all, MEs ‘brown’, ‘cyan’ and ‘darkturquoise’ showed the most significant positive correlations with temperature (Figure 2A and B). MEs ‘red and ‘grey60’ showed the most significant negative correlations with temperature (Figure 2A and B). Thus, MEs ‘brown’, ‘cyan’, ‘darkturquoise’, ‘red’, ‘grey60’ were identified as target modules (temperature responsive modules) for further analysis. The 2,985 genes in these modules were analyzed for their expression correlation with treatment temperature in DESeq2, and 2,856 genes with an FDR < 0.05 in the correlation were identified as target DEGs. Among these 2,856 target DEGs, 1,798 were positively correlated with temperature while 1,058 were negatively correlated with temperature. A full list of these temperature responsive genes is available in Supporting Material 2.

**Figure 2.**
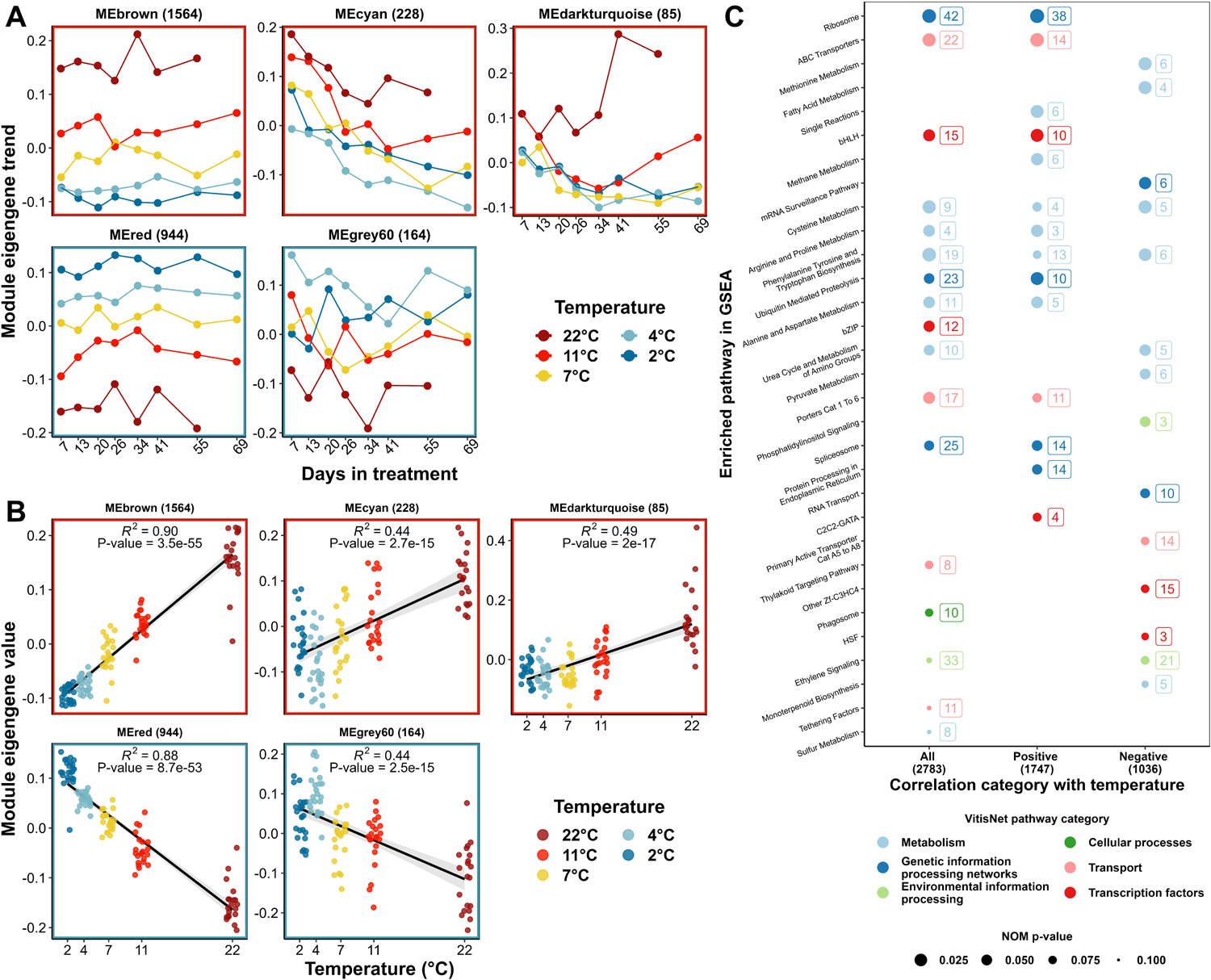
Identification of temperature responsive modules and pathway enrichment analysis of temperature responsive genes in Experiment 1. A) Model eigengenes (MEs) of temperature responsive modules observed across the experiment. B) Correlation analysis of the MEs of temperature responsive modules and treatment temperature; C) Pathway enrichment analysis of temperature responsive genes.

Target DEG were cross referenced with their corresponding CRIBI V1 annotations for pathway enrichment analysis. Three gene lists were generated based on their correlation categories with temperature (all, positive, or negative). Pre-ranked GSEA was separately conducted on three gene lists after ranking genes based on their correlation FDR. Results of GSEA are shown in Figure 2C. The number of significantly enriched pathways (NOM p-value < 0.1) was 17, 14 and 14 in the gene lists for all genes, the genes positively and negatively correlated to temperature, respectively (Figure 2C). These pathways comprise all major pathway classifications in VitisNet: Metabolism (12), Genetic information processing networks (6), Environmental information processing (2), Cellular processes (1), Transport (5) and Transcription factors (5) (Figure 2C).

The top 200 DEGs that showed significant negative correlation with temperature were used to identify transcriptomic response to low temperature in endodormant grapevine bud. Among these 200 DEGs, 147 were functionally annotated. Two pathways, phenylpropanoid biosynthesis (vv10940, VitisNet) and galactose metabolism (vv10052, VitisNet) were repeatedly represented among the top DEGs. The expressions of these DEGs in Experiment 1 and Experiment 2 are shown in Figure S6. We also investigated the top 200 DEGs that showed significant positive correlation with temperature, but no pathway was repeatedly represented among these DEGs.

### 3.4 Experiment 2: Differentiated phenotype but undifferentiated transcriptome

All buds remained dormant and unbroken during Experiment 2. The LTE of buds treated with stable temperatures of 11 °C, 7 °C and 2 °C remained unchanged within the range of −8 °C to −13 °C throughout the experiment (Figure 3A). In comparison, the LTE of buds treated with 7±5 °C in phase II diverged to a lower range of −12.5 °C to −18 °C during phase II regardless of their previous phase I treatment (Figure 3A). LTEs were also pooled based on phase II treatments (Figure 3B). The LTEs of phase II stable treatment and phase II 7±5 °C treatment show significant difference (p < 0.0001) in all the sample collections in phase II, indicating a differentiated phenotype of greater freezing tolerance during phase II. Pooled LTEs were also analyzed for their correlations with days in treatment in phase II (Figure 3C). The correlations are not significant, and the *R^2^*s are minimum (Figure 3C), indicating that freezing tolerance of pooled samples was not changed during phase II following the initial enhancement that occurred between 26 d and 34 d when samples were moved to the fluctuating temperature treatment.

**Figure 3.**
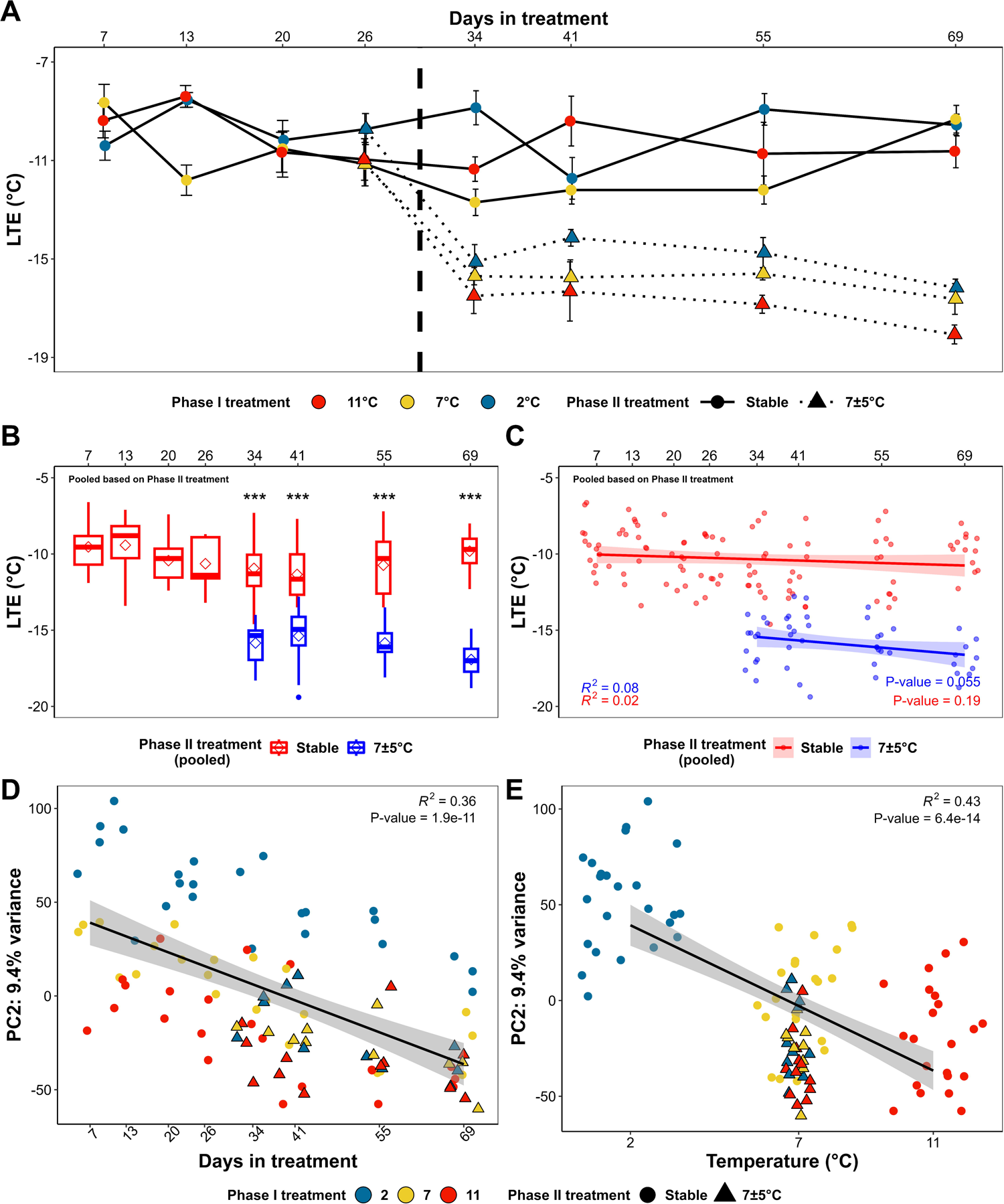
Freezing tolerance measurement and transcriptome characterization of grapevine buds in Experiment 2. A) Mean low temperature exotherm (LTE) observed across the experiment. Bars represent means ± SE (n = 5); B) Pooled LTE based on phase II treatment; C) Correlation analysis of pooled LTE and days in treatment; D) and E) Correlation analysis of PC2 with days in treatment and mean treatment temperature. Clustering of phase II samples and phase I 7 °C samples demonstrates convergence in the transcriptome of phase II treatment with phase I 7 °C.

PCA was applied on VST counts of all genes after low count filtering (16,028 genes) in Experiment 2 to characterize the transcriptome of the bud samples. PC1 and PC2 of all samples explain more than 27% of total variance, and the PC value of these two PCs are shown across the experiment in Figure S7. No apparent treatment effect can be identified in PC1 (Figure S7). Correlation analysis of PC2 and mean treatment temperature (mean treatment temperature of temperature cycling treatment in phase II is 7 °C) and days in treatment indicates that PC2 is not only negatively correlated with mean treatment temperature but also negatively correlated with days in treatment (Figure 3D and E).

The temperature responsive genes (2,856 genes) identified in Experiment 1 were also examined for their behavior in Experiment 2 to detect if these genes were differently regulated in the cycling treatment. PC1 and PC2 of all samples explain more than 35% of total variance (Figure S8A).

Correlation analysis of these PC values vs. mean treatment temperature and days in treatment demonstrates that PC1 of the samples correlated with mean treatment temperature, whereas PC2 of the samples significantly correlate with days in treatment (Figure S8B and C).

VST counts of detected genes (16,028 genes) after low count filtering in the temperature cycling experiment were subjected to WGCNA to construct gene co-expression network and detect gene co-expression modules. In total, 21 gene co-expression modules (excluding module ‘grey’) with number of genes ranging from 74 to 3,628 were identified. The MEs were tested for correlations with phase II treatment, days in treatment, and mean treatment temperature and were visualized across the experiment (Figure S9). Phase II treatment was transformed to a dummy variable (7±5 °C as ‘1’, stable as ‘0’) to facilitate correlation analysis. Although multiple MEs show significant correlations with days in treatment and mean treatment temperature, very few MEs (e.g., ME ‘black’, ‘royalblue’ and ‘greenyellow’) reveal significant correlations with phase II treatment, and the p-values of the significant correlations are high (Figure S9A). Based on ME visualization, the significance identified in these MEs-phase II treatment correlation analysis appear to be due to the segregation of the MEs of phase I 2 °C treatment samples from the other samples, or the time effect that differentiates phase I and phase II samples (Figure S9B). Thus, no target module specifically associated with the phenotypic shift in freezing tolerance was identified in the experiment. Instead, we conducted a contrast (stable vs. 7±5 °C) on all genes after low count filtering using DESeq2. This contrast resolved 87 genes with FDR < 0.05 and LFC > 2. The expressions of the functionally annotated genes among these genes are shown in Figure S10. Due in part to the limited number of DEGs in this contrast, pathway enrichment analysis did not identify any overrepresented pathways associated with the observed shift in freezing tolerance.

Further, we conducted three contrasts (phase I 2°C on 26 d vs. phase I 2°C × phase II 7±5°C on 34 d, phase I 7°C on 26 d vs. phase I 7°C × phase II 7±5°C on 34 d and phase I 11°C on 26 d vs. phase I 11°C × phase II 7±5°C on 34 d) to identify the shared DEGs in response to temperature cycling between 26 d and 34 d. No DEGs were identified after filtering with FDR< 0.05 and LFC >2.

Instead, filtering with FDR< 0.05 and LFC >1 identified 52 DEGs. The expression of the functional annotated genes among these 52 DEGs are shown in Figure S11.

## 4 Discussion

Although the leading threat posed to the survival of grapevine in winter is low temperatures that exceed the lethal threshold, an emerging issue is the prolonged period of warmth in fall because of climate change ^45,58^. As the first potential consequence, cold acclimation, which is progressively induced by low above and below freezing temperatures, may be compromised. Reduced acclimation in grapevine thus increases the susceptibility of grapevines to sudden temperature plunges ^17,59^. Secondly, as the accumulation of chilling is reduced under higher temperature in grapevine ^32,60^, prolonged maintenance of endodormancy under such temperatures might lead to unwanted physiological change of overwintering buds. However, a comprehensive examination of the physiology and transcriptome of endodormant grapevine under such temperature conditions have not been explored.

In this study, Experiment 1 examined the impact of different stable temperatures from 2 °C to 22 °C on the process of cold acclimation in grapevine endodormant buds. The treatment temperatures of 2 °C, 4 °C and 7 °C represent typical ambient temperatures that grapevine is exposed to under field conditions during the early stage of cold acclimation in fall, while the treatment temperatures of 11 °C and 22 °C partially simulate a prolonged period of warmth in the same window. Colder winters result in deeper freezing tolerance ^13,58,61,62^, thus we hypothesized that the growth chambers with low, steady temperatures (e.g., 2°C, 4°C) would gain freezing tolerance, relative to warm chambers. During the experiment, the LTEs of the buds in 22 °C, 11 °C, 7 °C and 2 °C did not show significant correlation with days in treatment (Figure 1B) while the LTEs of the buds in 4 °C showed significant negative correlation with time (Figure 1B). However, the apparent gain of freezing tolerance at 4 °C may be spurious given the lack of correlations in all of temperatures and particularly how there was no gain of phenotype at 2 °C. Thus, we conclude that the freezing tolerance of endodormant buds under all treatment temperatures remained unchanged and undifferentiated over the course of the experiment (Figure 1). In contrast to the unchanged phenotype, clear differences in the transcriptome were observed in the buds under different treatment temperatures with a total of 2,856 temperature responsive genes, comprising twenty-eight significantly enriched biological pathways (Figure 1 and 2).

Among the 2,856 temperature responsive genes, the expression of 1,798 genes was positively correlated with temperature, indicating the increase of transcript abundancy of these genes under higher temperatures. As a caveat of our sequencing approach, we are unable to conclusively determine if gene expression changes are solely to changes in transcription, rather than a result of other mechanisms such as the enhancement of mRNA stability or the decrease of mRNA decay ^63^. Assuming the increase is caused by upregulation, this pattern may indicate enhanced enzymatic activity under preferred temperature ^64^. However, since high temperature also potentially introduces pressure to avoid growth in endodormant buds, the mechanism of maintaining endodormancy under higher temperatures might be captured in this gene list. Pathway enrichment analysis of these genes revealed 14 significantly enriched pathways which are primarily associated with the categories of amino acid metabolism and genetic information processing. Many pathways in the major steps connecting DNA to mature proteins, splicing, translation and post-translation peptides modification were upregulated, suggesting the activation of some growth-related pathways as a potential ubiquitous response under growth permissive conditions. Interestingly, the ubiquitin-mediated proteolysis pathway, a major protein degradation mechanism in plants, was also found to be enriched. This proteolysis employs the binding of ubiquitin to target proteins through conjugating enzymes and the degradation of polyubiquitinated substrate by 26S proteasome complex ^65,66^. The upregulated genes in Experiment 1 include five genes encoding ubiquitin-conjugating enzyme E2 (*Vitvi02g00092*, *Vitvi08g00176*, *Vitvi08g01180*, *Vitvi15g01099* and *Vitvi16g01202*), two genes encoding seven in absentia protein (SINA) (*Vitvi14g01003* and *Vitvi15g01575*), one gene encoding mitogen-activated protein (MAP) kinase kinase (MEKK) (*Vitvi12g01775*), one gene encoding small ubiquitin-like modifier (SUMO) activating enzyme (*Vitvi03g00283*) and one gene encoding ubiquitin thioesterase (*Vitvi02g00247*). The upregulation of the nine genes (except for the ubiquitin thioesterase gene) would likely upregulate ubiquitin-mediated proteolysis pathway through promoting ubiquitin conjugation ^67^. The upregulation of ubiquitin-mediated proteolysis pathway has been reported as a response for protein turnover under various abiotic stresses ^67,68^ and the upregulation of this pathway has been observed in *Litchi chinensis* entering growth cessation and dormancy ^69^. Taking these results together, we propose a genetic control model of the maintenance of endodormancy under high temperatures (Figure 4). The enriched pathways include 1) spliceosome (vv23040, VitisNet); 2) ribosome (vv23010, VitisNet); 3) arginine and proline metabolism (vv10330, VitisNet), phenylalanine tyrosine and tryptophan biosynthesis (vv10400, VitisNet), cysteine metabolism (vv10272, VitisNet) and alanine and aspartate metabolism (vv10252, VitisNet); and 4) protein processing in endoplasmic reticulum (vv24141, VitisNet) and ubiquitin-mediated proteolysis (vv24120, VitisNet). Individual genes in these pathways are included in Supporting Material 2. Assuming that protein synthesis is unchanged, growth-related pathways are potentially activated under high temperatures, and proteins are potentially synthesized with a higher rate (compared to low temperatures) as a natural response to growth permissive conditions. The overproduction of proteins could be detected as a stress during endodormancy, inducing simultaneous upregulation of ubiquitin-mediated proteolysis, which in turn degrades extra proteins, thus maintaining dormancy (Figure 4). All the steps from pre-mRNA splicing to ubiquitin-mediated proteolysis require energy in the form of ATP ^70–73^. Our result and the proposed model suggest a hypothesis that endodormant buds are transcriptionally active and potentially metabolically active under growth permissive temperature, whereas the metabolites and proteins produced in this process may also be degraded, which could result in a waste of energy. The major energy source of dormant grapevine is carbohydrates, and the concentration of carbohydrates have been observed to correlate with freezing tolerance ^74–77^. While it has not yet been documented that carbohydrate level is necessary and sufficient to determine differences in supercooling ability in grapevine, it is possible that a waste of energy generated by carbohydrate catabolism during prolonged endodormancy under high temperatures in fall might compromise the ability of cold acclimation and threaten the survival of grapevine in winter. Future studies examining endodormant buds in warm temperatures, particularly those which examine the transcriptome, proteome, and metabolome in combination, are needed to help understand the apparent upregulation of genes involved in both protein synthesis and protein turnover observed in this study.

**Figure 4.**
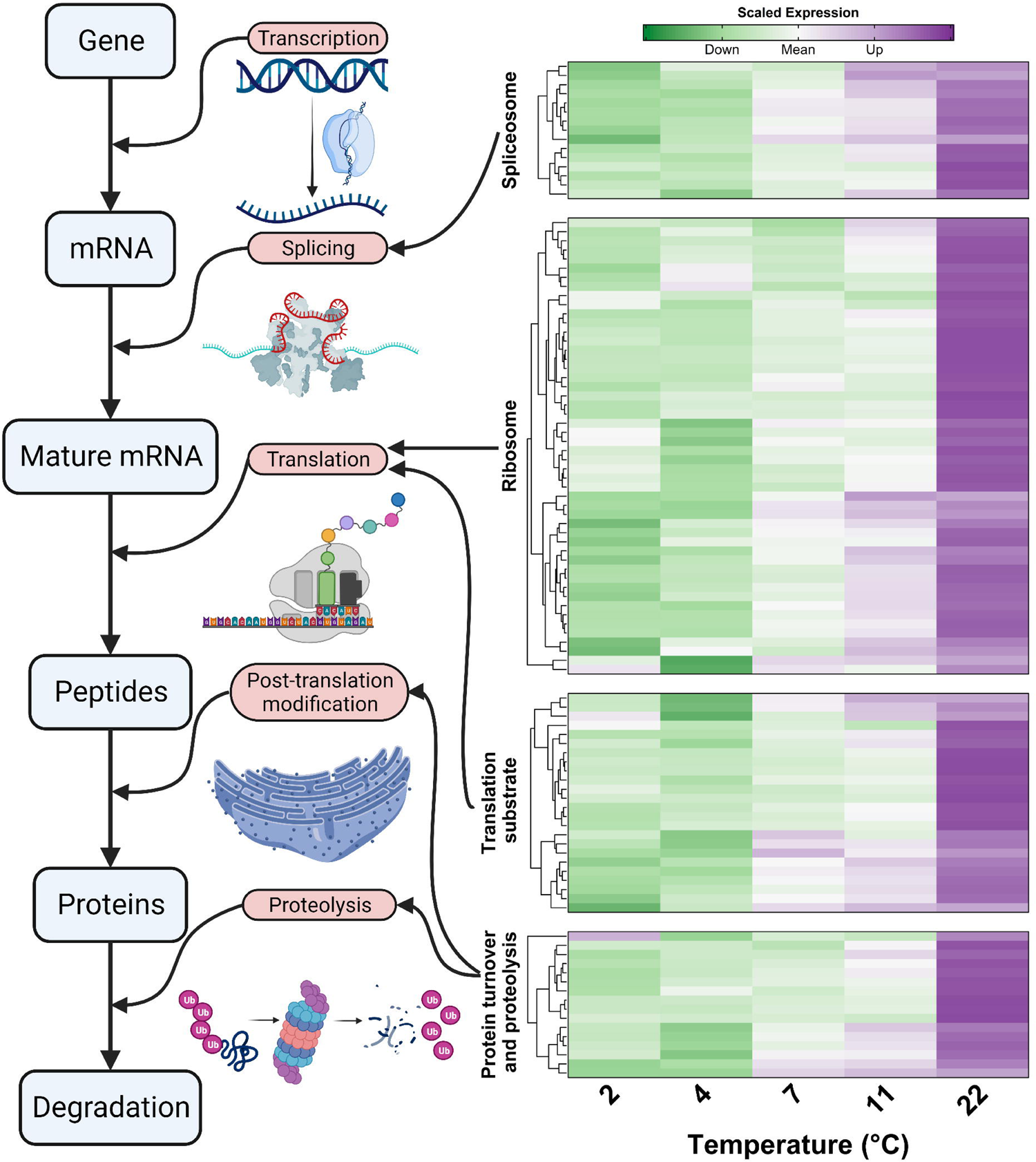
Schematic representation of a proposed model for the maintenance of endodormancy under high temperature conditions in grapevine. Blue boxes show the major substrates in genetic information processing and protein biosynthesis. Red boxes show the processes catalyzing the transitions between the substrates. The heatmap represents the DEGs in the enriched pathways in Experiment 1 that function during the transitions.

Among the 2,856 temperature responsive genes identified in Experiment 1, the expression of 1,058 genes were negatively correlated with temperature, indicating these genes are upregulated under lower temperatures. Although the increased expression of these genes did not result in the enhancement of freezing tolerance, these genes are likely a part of temperature sensing and cold response system in dormant grapevine buds. Among enriched pathways, the ethylene signaling pathway contained the greatest number of DEGs (Figure 2C). Ethylene has been reported for its importance in the control of freezing tolerance, however, the role of ethylene in the development of freezing tolerance and the response of ethylene signaling to low temperatures are inconsistent among different plant species ^22,78–82^. In our study, eight genes in the ethylene signaling pathway and 13 *ERF*s were upregulated under low temperature (Figure 5 and Figure S12). The DEGs in ethylene signaling include three genes (*Vitvi05g00684*, *Vitvi06g00518* and *Vitvi04g00115*) encoding ethylene receptors or co-receptors (two *ETR2* and one *RTE1*), four genes (*Vitvi05g00573*, *Vitvi18g00533*, *Vitvi18g00534* and *Vitvi11g00475*) encoding negative regulators (all three *CTR1* and one *EBF1*) and one gene (*Vitvi13g01126*) encoding transcription factor that stimulate the expression of *ERF* (*EIN3*) (Figure 5). Ethylene signaling has been linked to plant freezing tolerance through its regulation of *CBF/DREB* gene expression in various plant species ^79,82,83^. In absence of ethylene, ethylene receptors also inhibit ethylene signaling through promoting the activity of CTR1 ^84^, thus the upregulation of these genes most likely induce a repression of ethylene signaling. Despite significant changes in gene expression in this study, it seems the repression of ethylene signaling is not sufficient to alter the freezing tolerance phenotype of endodormant grapevine.

**Figure 5.**
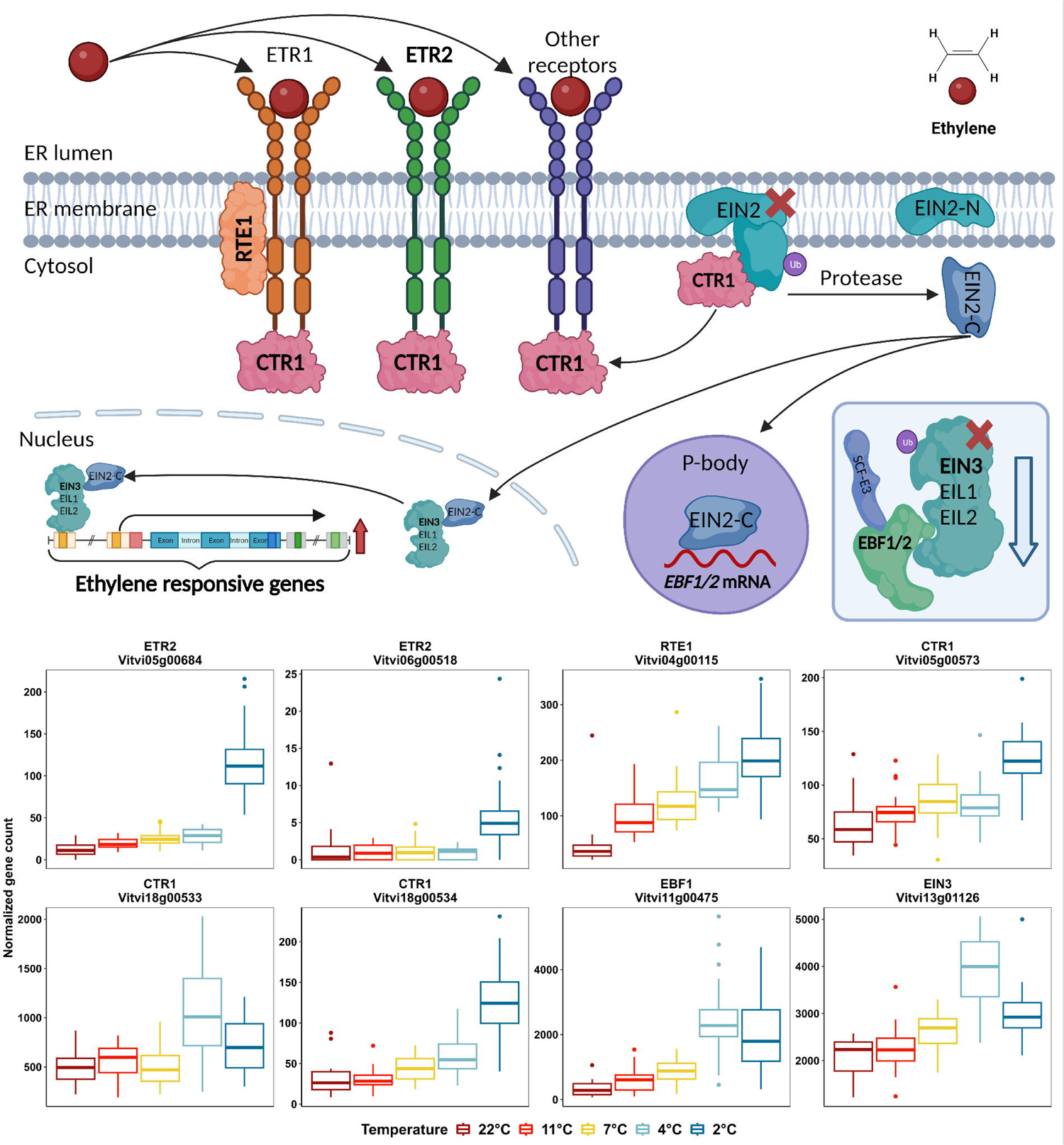
Impact of temperature on the DEGs in ethylene signaling pathway in Experiment 1. The schematic representation of ethylene signaling pathway in the figure shows a simplified ethylene signaling pathway that incorporated the major steps of the signaling and the proteins encoded by the DEGs identified in Experiment 1. Black arrows indicate the fate of the substrate when ethylene signaling is activated. Blue arrows and red arrows indicate the downregulation and the upregulation of the processes when ethylene signaling is activated, respectively. The protein encoded by the DEGs are bold-faced. Abbreviations: CTR, constitutive triple response protein; EBF, EIN3-binding F-box protein; EIL, EIN3-Like transcription factors; EIN, ethylene insensitive protein; ER, endoplasmic reticulum; ERF, ethylene response factors; ETR, ethylene receptors; P-body, processing body; RTE, reversion to ethylene sensitivity protein; SCF-E3, Skp1 Cullen F-box (SCF) E3 ubiquitin ligase complex; UB, ubiquitin.

The manual curation of the top 200 DEGs that negatively correlated with temperature in Experiment 1, two pathways that have been noted in previous cold related studies (galactose metabolism and phenylpropanoid biosynthesis) were identified (Figure S6A). In the galactose metabolism pathway, one gene encoding cell wall invertase (*Vitvi09g00193*), one gene encoding raffinose synthase (*Vitvi17g00885*) and one gene encoding stachyose synthase (*Vitvi07g00431*) exhibited higher expression in low temperature treatments (4 °C and 2 °C) (Figure S6A). Cell wall apoplastic invertase catalyzes the irreversible hydrolyzation of sucrose into glucose and sucrose, which are importance cryoprotectants for maintaining cell membrane integrity ^85,86^. Cell wall invertase genes were reported to be upregulated under cold stress in *Catharanthus roseus* ^87^. Raffinose synthase and stachyose synthase catalyze key reactions of the biosynthesis of raffinose and stachyose, two raffinose family oligosaccharides whose accumulation are correlated to the enhancement of low temperature tolerance in various plant species, including grapevine ^88–92^. In the phenylpropanoid biosynthesis pathway, low temperature responsive genes include a gene encoding cinnamoyl-CoA reductase (*Vitvi02g00717*), one gene encoding cinnamoyl alcohol dehydrogenase (*Vitvi13g02156*), one gene encoding resveratrol synthesis (*Vitvi16g01474*) and four genes encoding stilbene synthase (*Vitvi16g00988*, *Vitvi16g01470*, *Vitvi16g01474* and *Vitvi16g01484*) (Figure S6A). Stilbenes are a class of non-flavonoid polyphenols possessing a shared structure of two benzene rings connected by an ethylene bridge, and resveratrol is also a natural stilbene derivative found in grapevine and wine ^93,94^. These secondary metabolites are known as phytoalexins that are involved in the defensive responses to biotic and abiotic stresses ^94,95^. All of these five stilbene encoding genes (including the resveratrol encoding gene) were also found upregulated under freeze stress in grapevine single-bud green cuttings ^80^, suggesting that stilbenes may play a general role in cold temperature response in grapevine.

Experiment 2 examined the role of temperature fluctuation on the process of cold acclimation in endodormant buds. In contrast with Experiment 1, significant enhancement of freezing tolerance of buds was observed in the temperature cycling treatment in phase II regardless of the previous phase I treatment (Figure 3). Our first approach to identifying acclimation related gene expression involved contrasting the 2,856 temperature responsive genes identified in Experiment 1 to determine if their expression is impacted by the temperature cycling treatment. When the treatment was shifted from stable temperatures in phase I to cycling temperature in phase II, the expression of these genes in the 2 °C and 11 °C treatments converged on the expression profile observed in the stable 7 °C treatment (Figure S8). WGCNA analysis of only these temperature responsive genes was used to detect expression clustering and the major expression behavior in Experiment 2. Three co-expression modules were identified and the genes in the two major modules exhibited the converging behavior as shown in PCA (Figure S13). Similar behavior was also observed in most of the DEGs identified in Experiment 1 in the galactose metabolism pathway and phenylpropanoid biosynthesis pathway (Figure S6B). This result suggests that the temperature responsive genes identified in this study are likely associated with the mean temperature (mean temperature of temperature cycling treatment in phase II is 7 °C), but not daily temperature variation, and the alternation of the expression of these temperature responsive genes might not be correlated with the alternation of freezing tolerance. Neither PCA nor WGCNA identified DEGs specific to the shift in freezing tolerance observed in response to the phase II treatment. Instead, we contrasted stable versus cycling conditions (stable vs. 7±5 °C) for all genes and all samples to identify individual genes that respond to the temperature cycling effect. The contrast resulted in 87 DEGs, and 65 of these DEGs are functionally annotated in VitisNet. Based on the visualization for their expressions in the experiment (Figure S10). When examining the expression patterns of these genes, they fell into 4 different categories: 1) variation in expression related to the time effect that differentiates phase I samples and phase II samples (28 DEGs), 2) the temperature effects of stable 2 °C treatment (20 DEGs), 3) potential false positive results in lowly expressed genes that merely passed the low expression (13 DEGs) or 4) potential false positive results due to high expression variation (four DEGs) (Figure S10). The analysis of pooled LTE data indicates that the freezing tolerance of the buds under temperature cycling treatment was enhanced at the first timepoint in phase II (34 d after Experiment 2, corresponding to eight days under temperature cycling treatment) and maintained consistently lower than stable treatment but remained unchanged thereafter (Figure 3B and C). Assuming the enhanced freezing tolerance is a result of transcriptomic alternation, it is possible that mechanistic changes in gene expression needed for a change in freezing tolerance occurred within the eight days between the last timepoint in phase I (26 d) and the first timepoint in phase II (34 d). Using less stringent filtering criteria, 52 DEGs were identified though close examination of the patterns of expression change revealed only effects due to time, or false positive results from low gene counts or high replicate variation. Thus, we were unable to identify the putative mechanism for enhancement of freezing tolerance under the temperature cycling treatment using the contrasts. A likely explanation is that the promotive effect of temperature cycling on the freezing tolerance of endodormant bud is transient and was not captured with the temporal resolution of sample collections used in this study. Another, precisely designed experiment with higher sample collection, perhaps every 12 or 24 hours, is needed to uncover the transcriptomic control of the phenotypical change we observed in Experiment 2. Alternatively, the enhancement of freezing tolerance may result from translational, proteomic, lipidomic or fluidic response rather than an upstream transcriptomic response since the alternation of these mechanisms were also reported to affect plant freezing tolerance ^96–98^. To our knowledge, this study represents the first artificial induction of acclimation in grapevine, thus there is no established basis for the physiological mechanism for cold acclimation. Further investigation regarding these ‘omics’ needs to be conducted to better understand dormant grapevine bud’s response to temperature cycling.

Concerning the *ICE-CBF-COR* cascade, known for its regulatory and functional role in enhancing freezing tolerance under low temperature in plants, we did not identify any treatment response in the expression of any ICE, DREB or CBF genes in both experiments presented here. Among all the 14 ICE, CBF/DREB genes annotated in VitisNet, six genes passed the low count filtering. None of these six genes are CBF genes, and their expression patterns do not correlate with any treatment effect in our experiments (Figure S14 and S15). This finding does not agree with many previous studies of grapevine reporting the upregulation of CBF genes in response to low temperature treatment and the enhancement of freezing tolerance under stable low temperature treatment ^99–104^. Although the undetectable expression of CBF genes might be a result of the lower-depth sequencing method used in our study, another likely and plausible reason is that the experimental material used in previous studies are either green tissues (leaves, shoot tips or paradormant buds) in actively growing grapevines or overexpressing lines of these genes in *A. thaliana*, whereas we used endodormant single-bud cuttings. Consequently, the findings in previous studies may be less appropriate for understanding how grapevine initiates cold acclimation during endodormancy. Although the function of *ICE-CBF-COR* cascade has been intensive studied over the last two decades, findings are mainly based on annual model plants ^82,105–108^. The understanding of *ICE-CBF-COR* cascade regulation in the cold acclimation of dormant woody perennials is still limited ^109^. Our result suggests that the current understanding *ICE-CBF-COR* cascade should not be generalized between plant species and even between different phenological stages of a same species. Further precise studies need to be conducted to elucidate the functionality of *ICE-CBF-COR* cascade in cold acclimation of endodormant grapevine.

In this study we examined freezing tolerance and transcriptomics of endodormant grapevine buds and the impact on cold acclimation under different temperature regimes. Our result shows that non-freezing stable temperatures did not alter the freezing tolerance in endodormant buds, but temperature cycling seems essential for the enhancement of freezing tolerance. In Experiment 1, we identified a list of genes that are highly expressed under high stable temperatures and proposed a proteolysis-mediated mechanistic model for the maintenance of endodormancy under high temperature conditions in grapevine. We also identified a list of genes that are highly expressed under low stable temperatures and detected potential alternations of several previously reported pathways that function in freezing tolerance, including the ethylene signaling pathway, the galactose metabolism pathway and the phenylpropanoid biosynthesis pathway. However, the alternation of these pathways was not shown to drive changes in freezing tolerance. In Experiment 2, we successfully promoted acclimation in endodormant grapevine buds, resulting in an increase and subsequent maintenance of 5 °C of freezing tolerance. This result leads us to conclude that the process of cold acclimation in endodormant grapevine requires temperature fluctuation to effectively drive changes in freezing tolerance. This result is perhaps not surprising given the natural diurnal fluctuation of temperature that grapevines are exposed to in field conditions. It is however surprising to note that the movement of all three stable temperature treatments into a shared cycling treatment resulted in comparably similar freezing tolerance. This result suggests that the cycle we used induces a specific maximum depth of freezing tolerance and this information could be valuable for the modeling of freezing tolerance development of grapevines. The effects of other temperature cycles on the acclimation process, both in amplitude and frequency, remain to be studied. In contrast to the success in shifting the phenotype of cold acclimation. Our analysis of the transcriptomic response was inconclusive, presumably due to the resolution of sample collection. Manual examination of the genes encoding for ICE or CBF/DREB indicates that these genes did not show differential expression under different temperature regimes, and their expression did not correlate with freezing tolerance. While the results of this study shed light on the process of cold acclimation in grapevine and demonstrate that temperature cycling plays a critical role, future, high-resolution studies are needed to directly link transcriptomic and other physiological changes and are necessary for the development of mitigation methods to enhance cold acclimation as climate changes.

## Supporting information

Fig. S1-S15 will be used for the link to the file on the preprint site

Supporting Material 2 will be used for the link to the file on the preprint site

## 5 Acknowledgement

This study was supported in part by the USDA -ARS appropriated project 1910–21220–006–00D and by the National Institute of Food and Agriculture, U.S. Department of Agriculture, through the Northeast Sustainable Agriculture Research and Education program under subaward number GNE16-130. We would like to acknowledge Hanna Martens, Amy Swezc-McFadden, and Kathleen Deys for assistance in collecting dormant cuttings and in processing samples for LTE analysis.

## DATA AVAILABILITY STATEMENT

All RNA-seq raw data along with processed gene count matrix and sample metadata are available in NCBI-GEO (accession: GSE232062).

## Notes

https://www.ncbi.xyz/geo/query/acc.cgi?acc=GSE232062

### Competing Interest Statement

The authors have declared no competing interest.

## References

1 Boadu FO. Chapter 17 - Climate Change. In: Boadu FO (ed). Agricultural Law and Economics in Sub-Saharan Africa. Academic Press: San Diego, 2016, pp 555–571.

2 Raza A, Razzaq A, Mehmood SS, et al. Impact of Climate Change on Crops Adaptation and Strategies to Tackle Its Outcome: A Review. Plants 2019; 8: 34.

3 Salama A-M, Ezzat A, El-Ramady H, et al. Temperate Fruit Trees under Climate Change: Challenges for Dormancy and Chilling Requirements in Warm Winter Regions. Horticulturae 2021; 7: 86.

4 Kling GW, Hayhoe K, Johnson LB et al. Confronting climate change in the Great Lakes region: impacts on our communities and ecosystems. Union of Concerned Scientists; Ecological Society of America, 2003.

5 Hong C, Mueller ND, Burney JA et al. Impacts of ozone and climate change on yields of perennial crops in California. Nat Food 2020; 1: 166–172.

6 Leisner CP. Review: Climate change impacts on food security-focus on perennial cropping systems and nutritional value. Plant Sci 2020; 293: 110412.

7 Schnabel BJ, Wample RL. Dormancy and Cold Hardiness in Vitis vinifera L. cv. White Riesling as Influenced by Photoperiod and Temperature. Am J Enol Vitic 1987; 38: 265–272.

8 White MA, Diffenbaugh NS, Jones GV, Pal JS, Giorgi F. Extreme heat reduces and shifts United States premium wine production in the 21st century. Proc Natl Acad Sci 2006; 103: 11217–11222.

9 Molitor D, Caffarra A, Sinigoj P, Pertot I, Hoffmann L, Junk J. Late frost damage risk for viticulture under future climate conditions: a case study for the Luxembourgish winegrowing region: Late frost damage risk. Aust J Grape Wine Res 2014; 20: 160–168.

10 Mozell MR, Thach L. The impact of climate change on the global wine industry: Challenges & solutions. Wine Econ Policy 2014; 3: 81–89.

11 Mosedale JR, Abernethy KE, Smart RE, Wilson RJ, Maclean IMD. Climate change impacts and adaptive strategies: lessons from the grapevine. Glob Change Biol 2016; 22: 3814–3828.

12 Droulia F, Charalampopoulos I. Future Climate Change Impacts on European Viticulture: A Review on Recent Scientific Advances. Atmosphere 2021; 12: 495.

13 Zabadal TJ, Dami IE, Goffinet MC, Martinson TE, Chien ML. Winter injury to grapevines and methods of protection. Michigan State University Extension, 2007.

14 Wake CMF, Fennell A. Morphological, physiological and dormancy responses of three Vitis genotypes to short photoperiod. Physiol Plant 2000; 109: 203–210.

15 Gusta LV, Trischuk R, Weiser CJ. Plant Cold Acclimation: The Role of Abscisic Acid. J Plant Growth Regul 2005; 24: 308–318.

16 Thomashow MF. PLANT COLD ACCLIMATION: Freezing Tolerance Genes and Regulatory Mechanisms. Annu Rev Plant Physiol Plant Mol Biol 1999; 50: 571–599.

17 Hincha DK, Zuther E. Introduction: Plant Cold Acclimation and Winter Survival. In: Hincha DK, Zuther E (eds). Plant Cold Acclimation: Methods and Protocols. Springer US: New York, NY, 2020, pp 1–7.

18 Hu Y, Jiang L, Wang F, Yu D. Jasmonate Regulates the INDUCER OF CBF EXPRESSION-C-REPEAT BINDING FACTOR/DRE BINDING FACTOR1 Cascade and Freezing Tolerance in Arabidopsis. Plant Cell 2013; 25: 2907–2924.

19 Jiang B, Shi Y, Zhang X, et al. PIF3 is a negative regulator of the CBF pathway and freezing tolerance in Arabidopsis. Proc Natl Acad Sci 2017; 114: E6695–E6702.

20 Ding Y, Shi Y, Yang S. Advances and challenges in uncovering cold tolerance regulatory mechanisms in plants. New Phytol 2019; 222: 1690–1704.

21 Rubio S, Pérez FJ. ABA and its signaling pathway are involved in the cold acclimation and deacclimation of grapevine buds. Sci Hortic 2019; 256: 108565.

22 Shi Y, Tian S, Hou L et al. Ethylene Signaling Negatively Regulates Freezing Tolerance by Repressing Expression of CBF and Type-A ARR Genes in Arabidopsis. Plant Cell 2012; 24: 2578–2595.

23 Guy C, Kaplan F, Kopka J, Selbig J, Hincha DK. Metabolomics of temperature stress. Physiol Plant 2008; 132: 220–235.

24 Korn M, Peterek S, Mock H-P, Heyer AG, Hincha DK. Heterosis in the freezing tolerance, and sugar and flavonoid contents of crosses between Arabidopsis thaliana accessions of widely varying freezing tolerance. Plant Cell Environ 2008; 31: 813–827.

25 Bredow M, Walker VK. Ice-Binding Proteins in Plants. Front Plant Sci 2017; 8: 2153.

26 Gusta LV, Wisniewski M. Understanding plant cold hardiness: an opinion. Physiol Plant 2013; 147: 4–14.

27 Arias NS, Bucci SJ, Scholz FG, Goldstein G. Freezing avoidance by supercooling in Olea europaea cultivars: the role of apoplastic water, solute content and cell wall rigidity. Plant Cell Environ 2015; 38: 2061–2070.

28 Ritonga FN, Chen S. Physiological and Molecular Mechanism Involved in Cold Stress Tolerance in Plants. Plants 2020; 9: 560.

29 Lang GA. Dormancy: a new universal terminology. HortScience 1987; 22: 817–822.

30 Campoy JA, Ruiz D, Egea J. Dormancy in temperate fruit trees in a global warming context: A review. Sci Hortic 2011; 130: 357–372.

31 Kovaleski AP. Woody species do not differ in dormancy progression: Differences in time to budbreak due to forcing and cold hardiness. Proc Natl Acad Sci 2022; 119: e2112250119.

32 Londo JP, Johnson LM. Variation in the chilling requirement and budburst rate of wild Vitis species. Environ Exp Bot 2014; 106: 138–147.

33 Bielenberg DG, Wang Y (Eileen), Li Z et al. Sequencing and annotation of the evergrowing locus in peach [Prunus persica (L.) Batsch] reveals a cluster of six MADS-box transcription factors as candidate genes for regulation of terminal bud formation. Tree Genet Genomes 2008; 4: 495–507.

34 Li Z, Reighard GL, Abbott AG, Bielenberg DG. Dormancy-associated MADS genes from the EVG locus of peach [Prunus persica (L.) Batsch] have distinct seasonal and photoperiodic expression patterns. J Exp Bot 2009; 60: 3521–3530.

35 Yamane H, Ooka T, Jotatsu H, Hosaka Y, Sasaki R, Tao R. Expressional regulation of PpDAM5 and PpDAM6, peach (Prunus persica) dormancy-associated MADS-box genes, by low temperature and dormancy-breaking reagent treatment. J Exp Bot 2011; 62: 3481–3488.

36 Yu J, Conrad AO, Decroocq V et al. Distinctive Gene Expression Patterns Define Endodormancy to Ecodormancy Transition in Apricot and Peach. Front Plant Sci 2020; 11: 180.

37 Sasaki R, Yamane H, Ooka T et al. Functional and Expressional Analyses of PmDAM Genes Associated with Endodormancy in Japanese Apricot. Plant Physiol 2011; 157: 485–497.

38 Tuan PA, Bai S, Saito T, Ito A, Moriguchi T. Dormancy-Associated MADS-Box (DAM) and the Abscisic Acid Pathway Regulate Pear Endodormancy Through a Feedback Mechanism. Plant Cell Physiol 2017; 58: 1378–1390.

39 Wang J, Gao Z, Li H et al. Dormancy-Associated MADS-Box (DAM) Genes Influence Chilling Requirement of Sweet Cherries and Co-Regulate Flower Development with SOC1 Gene. Int J Mol Sci 2020; 21: 921.

40 Niu Q, Li J, Cai D et al. Dormancy-associated MADS-box genes and microRNAs jointly control dormancy transition in pear (Pyrus pyrifolia white pear group) flower bud. J Exp Bot 2016; 67: 239–257.

41 Quesada-Traver C, Guerrero BI, Badenes ML, Rodrigo J, Ríos G, Lloret A. Structure and Expression of Bud Dormancy-Associated MADS-Box Genes (DAM) in European Plum. Front Plant Sci 2020; 11: 1288.

42 Rothkegel K, Sandoval P, Soto E et al. Dormant but Active: Chilling Accumulation Modulates the Epigenome and Transcriptome of Prunus avium During Bud Dormancy. Front Plant Sci 2020; 11: 1115.

43 Kamal N, Ochßner I, Schwandner A et al. Characterization of genes and alleles involved in the control of flowering time in grapevine. PLOS ONE 2019; 14: e0214703.

44 Luedeling E, Girvetz EH, Semenov MA, Brown PH. Climate Change Affects Winter Chill for Temperate Fruit and Nut Trees. PLoS ONE 2011; 6: e20155.

45 Luedeling E. Climate change impacts on winter chill for temperate fruit and nut production: A review. Sci Hortic 2012; 144: 218–229.

46 Eichhorn KW, Lorenz DH. Phenological development stages of the grapevine. Nachrichtenblatt Dtsch Pflanzenschutzdienstes 1977; 29: 119–120.

47 Mills LJ, Ferguson JC, Keller M. Cold-Hardiness Evaluation of Grapevine Buds and Cane Tissues. Am J Enol Vitic 2006; 57: 194–200.

48 Londo JP, Moyer MM, Mireles M et al. Evaluation of Sample Preparation Practices Common with Differential Thermal Analysis of Grapevine Bud Cold Hardiness. Am J Enol Vitic 2023; 74: 0740002.

49 Pierquet P, Stushnoff C. Relationship of Low Temperature Exotherms to Cold Injury in Vitis Riparia Michx. Am J Enol Vitic 1980; 31: 1–6.

50 Andrew S. FastQC. 2010.http://www.bioinformatics.babraham.ac.uk/projects/fastqc/.

51 Nordberg H, Cantor M, Dusheyko S et al. The genome portal of the Department of Energy Joint Genome Institute: 2014 updates. Nucleic Acids Res 2014; 42: D26–D31.

52 Dobin A, Davis CA, Schlesinger F et al. STAR: ultrafast universal RNA-seq aligner. Bioinformatics 2013; 29: 15–21.

53 Canaguier A, Grimplet J, Di Gaspero G et al. A new version of the grapevine reference genome assembly (12X.v2) and of its annotation (VCost.v3). Genomics Data 2017; 14: 56–62.

54 Love MI, Huber W, Anders S. Moderated estimation of fold change and dispersion for RNA-seq data with DESeq2. Genome Biol 2014; 15: 550.

55 Langfelder P, Horvath S. WGCNA: an R package for weighted correlation network analysis. BMC Bioinformatics 2008; 9: 559.

56 Subramanian A, Tamayo P, Mootha VK et al. Gene set enrichment analysis: A knowledge-based approach for interpreting genome-wide expression profiles. Proc Natl Acad Sci 2005; 102: 15545–15550.

57 Grimplet J, Cramer GR, Dickerson JA, Mathiason K, Van Hemert J, Fennell AY. VitisNet: “Omics” Integration through Grapevine Molecular Networks. PLOS ONE 2009; 4: e8365.

58 De Rosa V, Vizzotto G, Falchi R. Cold Hardiness Dynamics and Spring Phenology: Climate-Driven Changes and New Molecular Insights Into Grapevine Adaptive Potential. Front Plant Sci 2021; 12: 591.

59 Howell GS. Grapevine cold hardiness: Mechanisms of cold acclimation, mid-winter hardiness maintenance, and spring deacclimation. Amer J Enol Viticult 2000; 51: 35–48.

60 Dokoozlian NK. Chilling Temperature and Duration Interact on the Budbreak of ‘Perlette’ Grapevine Cuttings. HortScience 1999; 34: 1–3.

61 Davenport JR, Keller M, Mills LJ. How Cold Can You Go? Frost and Winter Protection for Grape. HortScience 2008; 43: 1966–1969.

62 Londo JP, Kovaleski AP. Deconstructing cold hardiness: variation in supercooling ability and chilling requirements in the wild grapevine *Vitis riparia*: Cold hardiness in *Vitis riparia*. Aust J Grape Wine Res 2019; 25: 276–285.

63 Agarwal V, Kelley DR. The genetic and biochemical determinants of mRNA degradation rates in mammals. Genome Biol 2022; 23: 245.

64 Ding Y, Shi Y, Yang S. Molecular Regulation of Plant Responses to Environmental Temperatures. Mol Plant 2020; 13: 544–564.

65 Ciechanover A, Orian A, Schwartz AL. Ubiquitin-mediated proteolysis: biological regulation via destruction. BioEssays 2000; 22: 442–451.

66 Moon J, Parry G, Estelle M. The Ubiquitin-Proteasome Pathway and Plant Development. Plant Cell 2004; 16: 3181–3195.

67 Sharma B, Joshi D, Yadav PK, Gupta AK, Bhatt TK. Role of Ubiquitin-Mediated Degradation System in Plant Biology. Front Plant Sci 2016; 7: 806.

68 Doroodian P, Hua Z. The Ubiquitin Switch in Plant Stress Response. Plants 2021; 10: 246.

69 Zhang H, Li H, Lai B, Xia H, Wang H, Huang X. Morphological Characterization and Gene Expression Profiling during Bud Development in a Tropical Perennial, Litchi chinensis Sonn. Front Plant Sci 2016; 7: 1517.

70 Peth A, Nathan JA, Goldberg AL. The ATP Costs and Time Required to Degrade Ubiquitinated Proteins by the 26 S Proteasome. J Biol Chem 2013; 288: 29215–29222.

71 Lynch M, Marinov GK. The bioenergetic costs of a gene. Proc Natl Acad Sci U S A 2015; 112: 15690–15695.

72 Tseng C-K, Chung C-S, Chen H-C, Cheng S-C. A central role of Cwc25 in spliceosome dynamics during the catalytic phase of pre-mRNA splicing. RNA 2017; 23: 546–556.

73 Martinez-Seidel F, Beine-Golovchuk O, Hsieh Y-C, Kopka J. Systematic Review of Plant Ribosome Heterogeneity and Specialization. Front Plant Sci 2020; 11: 948.

74 Grant TN, Dami IE, Ji T, Scurlock D, Streeter J. Variation in leaf and bud soluble sugar concentration among Vitis genotypes grown under two temperature regimes. Can J Plant Sci 2009; 89: 961–968.

75 Grant TNL, Dami IE. Physiological and Biochemical Seasonal Changes in Vitis Genotypes with Contrasting Freezing Tolerance. Am J Enol Vitic 2015; 66: 195–203.

76 Wang H, Dami IE. Evaluation of Budbreak-Delaying Products to Avoid Spring Frost Injury in Grapevines. Am J Enol Vitic 2020; 71: 181–190.

77 De Rosa V, Falchi R, Moret E, Vizzotto G. Insight into Carbohydrate Metabolism and Signaling in Grapevine Buds during Dormancy Progression. Plants 2022; 11: 1027.

78 Catalá R, Salinas J. The Arabidopsis ethylene overproducer mutant eto1-3 displays enhanced freezing tolerance. Plant Signal Behav 2015; 10: e989768.

79 Sun X, Zhao T, Gan S et al. Ethylene positively regulates cold tolerance in grapevine by modulating the expression of ETHYLENE RESPONSE FACTOR 057. Sci Rep 2016; 6: 24066.

80 Londo JP, Kovaleski AP, Lillis JA. Divergence in the transcriptional landscape between low temperature and freeze shock in cultivated grapevine (Vitis vinifera). Hortic Res 2018; 5: 1–14.

81 Wang Y, Jiang H, Mao Z et al. Ethylene increases the cold tolerance of apple via the MdERF1B–MdCIbHLH1 regulatory module. Plant J 2021; 106: 379–393.

82 Hwarari D, Guan Y, Ahmad B et al. ICE-CBF-COR Signaling Cascade and Its Regulation in Plants Responding to Cold Stress. Int J Mol Sci 2022; 23: 1549.

83 Robison JD, Yamasaki Y, Randall SK. The Ethylene Signaling Pathway Negatively Impacts CBF/DREB-Regulated Cold Response in Soybean (Glycine max). Front Plant Sci 2019; 10: 121.

84 Binder BM. Ethylene signaling in plants. J Biol Chem 2020; 295: 7710–7725.

85 Ma Y, Zhang Y, Lu J, Shao H. Roles of plant soluble sugars and their responses to plant cold stress. Afr J Biotechnol 2009; 8: 2004–2010.

86 Proels RK, Hückelhoven R. Cell_Jwall invertases, key enzymes in the modulation of plant metabolism during defence responses. Mol Plant Pathol 2014; 15: 858–864.

87 Nishanth MJ, Sheshadri SA, Rathore SS, Srinidhi S, Simon B. Expression analysis of Cell wall invertase under abiotic stress conditions influencing specialized metabolism in Catharanthus roseus. Sci Rep 2018; 8: 15059.

88 Sun X, Matus JT, Wong DCJ et al. The GARP/MYB-related grape transcription factor AQUILO improves cold tolerance and promotes the accumulation of raffinose family oligosaccharides. J Exp Bot 2018; 69: 1749–1764.

89 Han Q, Qi J, Hao G et al. ZmDREB1A Regulates RAFFINOSE SYNTHASE Controlling Raffinose Accumulation and Plant Chilling Stress Tolerance in Maize. Plant Cell Physiol 2020; 61: 331–341.

90 Wang H, Blakeslee JJ, Jones ML, Chapin LJ, Dami IE. Exogenous abscisic acid enhances physiological, metabolic, and transcriptional cold acclimation responses in greenhouse-grown grapevines. Plant Sci 2020; 293: 110437.

91 Khan M, Hu J, Dahro B et al. ERF108 from Poncirus trifoliata (L.) Raf. functions in cold tolerance by modulating raffinose synthesis through transcriptional regulation of PtrRafS. Plant J 2021; 108: 705–724.

92 Sanyal R, Kumar S, Pattanayak A, Kar A, Bishi SK. Optimizing raffinose family oligosaccharides content in plants: A tightrope walk. Front Plant Sci 2023; 14: 1134754.

93 Bertelli AA, Ferrara F, Diana G et al. Resveratrol, a natural stilbene in grapes and wine, enhances intraphagocytosis in human promonocytes: a co-factor in antiinflammatory and anticancer chemopreventive activity. Int J Tissue React 1999; 21: 93–104.

94 Valletta A, Iozia LM, Leonelli F. Impact of Environmental Factors on Stilbene Biosynthesis. Plants 2021; 10: 90.

95 Qayyum Z, Noureen F, Khan M, et al. Identification and Expression Analysis of Stilbene Synthase Genes in Arachis hypogaea in Response to Methyl Jasmonate and Salicylic Acid Induction. Plants 2022; 11: 1776.

96 Takahashi D, Li B, Nakayama T, Kawamura Y, Uemura M. Plant plasma membrane proteomics for improving cold tolerance. Front Plant Sci 2013; 4: 90.

97 Song Y, Vu HS, Shiva S et al. A lipidomic approach to identify cold-induced changes in Arabidopsis membrane lipid composition. Methods Mol Biol Clifton NJ 2020; 2156: 187–202.

98 Cano-Ramirez DL, Carmona-Salazar L, Morales-Cedillo F, Ramírez-Salcedo J, Cahoon EB, Gavilanes-Ruíz M. Plasma Membrane Fluidity: An Environment Thermal Detector in Plants. Cells 2021; 10: 2778.

99 Xiao H, Siddiqua M, Braybrook S, Nassuth A. Three grape CBF/DREB1 genes respond to low temperature, drought and abscisic acid. Plant Cell Environ 2006; 29: 1410–1421.

100 Xiao H, Tattersall E a. R, Siddiqua MK, Cramer GR, Nassuth A. CBF4 is a unique member of the CBF transcription factor family of Vitis vinifera and Vitis riparia. Plant Cell Environ 2008; 31: 1–10.

101 Li J, Wang L, Zhu W, Wang N, Xin H, Li S. Characterization of two VvICE1 genes isolated from ‘Muscat Hamburg’ grapevine and their effect on the tolerance to abiotic stresses. Sci Hortic 2014; 165: 266–273.

102 Karimi M, Ebadi A, Mousavi SA, Salami SA, Zarei A. Comparison of CBF1, CBF2, CBF3 and CBF4 expression in some grapevine cultivars and species under cold stress. Sci Hortic 2015; 197: 521–526.

103 Rubio S, Noriega X, Pérez FJ. Abscisic acid (ABA) and low temperatures synergistically increase the expression of CBF/DREB1 transcription factors and cold-hardiness in grapevine dormant buds. Ann Bot 2019; 123: 681–689.

104 Wang L, Sadeghnezhad E, Riemann M, Nick P. Microtubule dynamics modulate sensing during cold acclimation in grapevine suspension cells. Plant Sci 2019; 280: 18–30.

105 Cook D, Fowler S, Fiehn O, Thomashow MF. A prominent role for the CBF cold response pathway in configuring the low-temperature metabolome of Arabidopsis. Proc Natl Acad Sci U S A 2004; 101: 15243–15248.

106 Shi Y, Huang J, Sun T et al. The precise regulation of different COR genes by individual CBF transcription factors in Arabidopsis thaliana. J Integr Plant Biol 2017; 59: 118–133.

107 Li Y, Wang X, Ban Q et al. Comparative transcriptomic analysis reveals gene expression associated with cold adaptation in the tea plant Camellia sinensis. BMC Genomics 2019; 20: 624.

108 Liu Y, Dang P, Liu L, He C. Cold acclimation by the CBF–COR pathway in a changing climate: Lessons from Arabidopsis thaliana. Plant Cell Rep 2019; 38: 511–519.

109 Wisniewski M, Nassuth A, Arora R. Cold Hardiness in Trees: A Mini-Review. Front Plant Sci 2018; 9: 1394.

