## Supplementary material for "Physiological and transcriptomic characterization of cold acclimation in endodormant grapevine under different temperature regimes": Fig. S1-S15 will be used for the link to the file on the preprint site

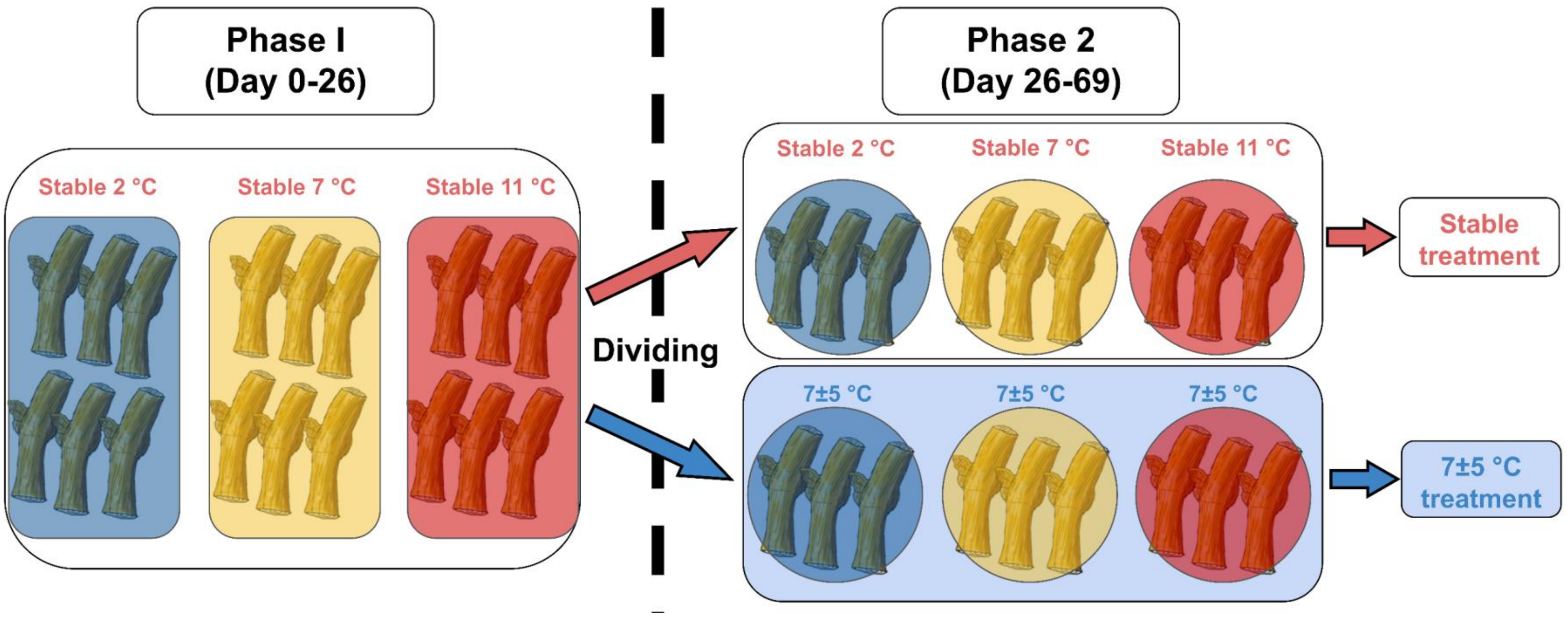


**Figure S1. Schematic presentation of experimental design of Experiment 2.** During phase I (0 d to 26 d), three stable temperature treatments (2 °C, 7 °C and 11 °C) were applied on the single-bud cutting in three growth chambers. During phase II (26 d to d 69 d), half of the cuttings in each growth chamber were moved to another growth chamber to be subjected to 7±5 °C temperature cycling treatment, and the remaining halves were subjected to the same treatments in phase I.

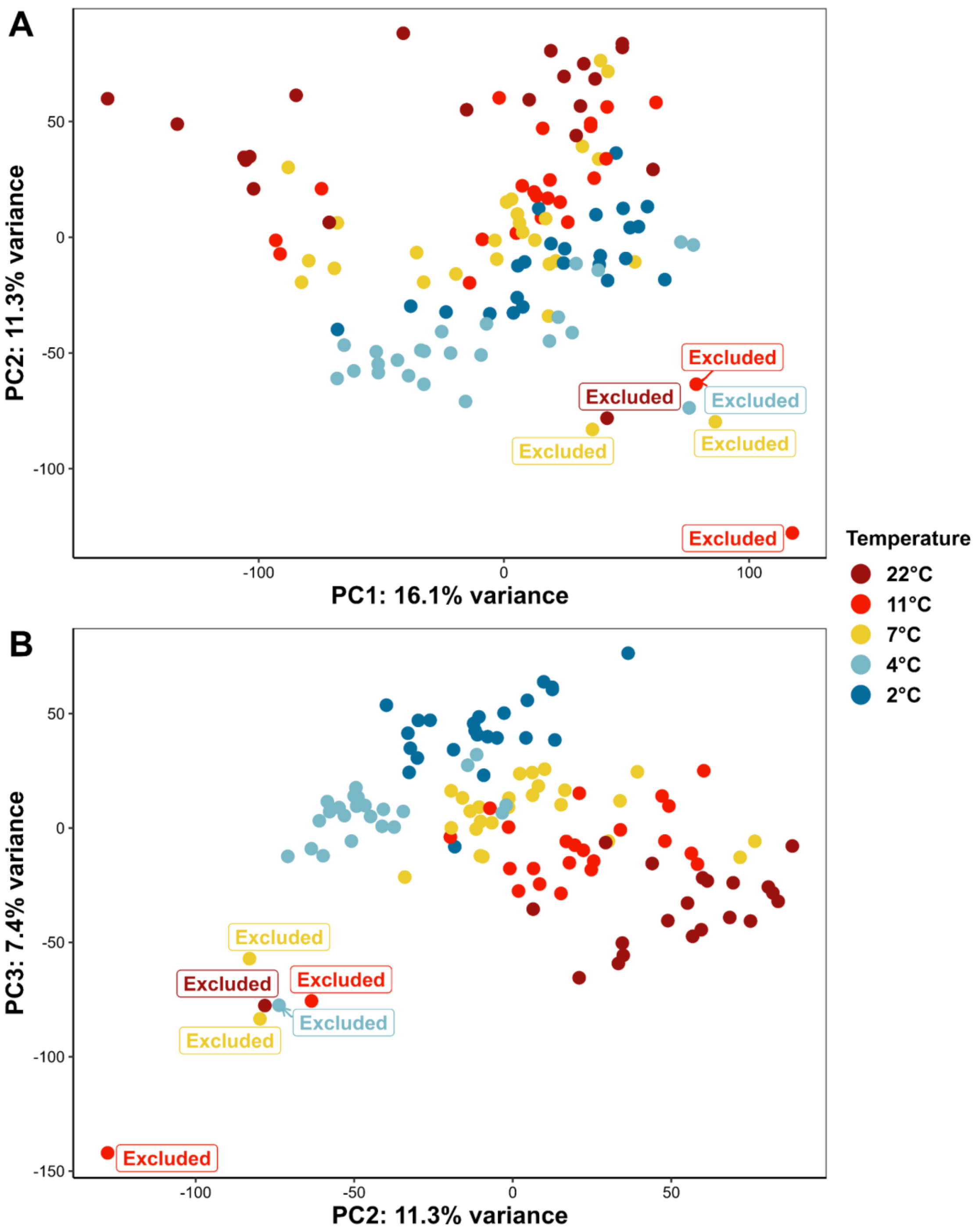


**Figure S2. Principal component analysis (PCA) of RNA-seq samples in Experiment 1 for outlier filtering.** A) principal component one (PC1) and PC2; B) PC2 and PC3. The six samples that derived from the main cluster in all three PCs were excluded.

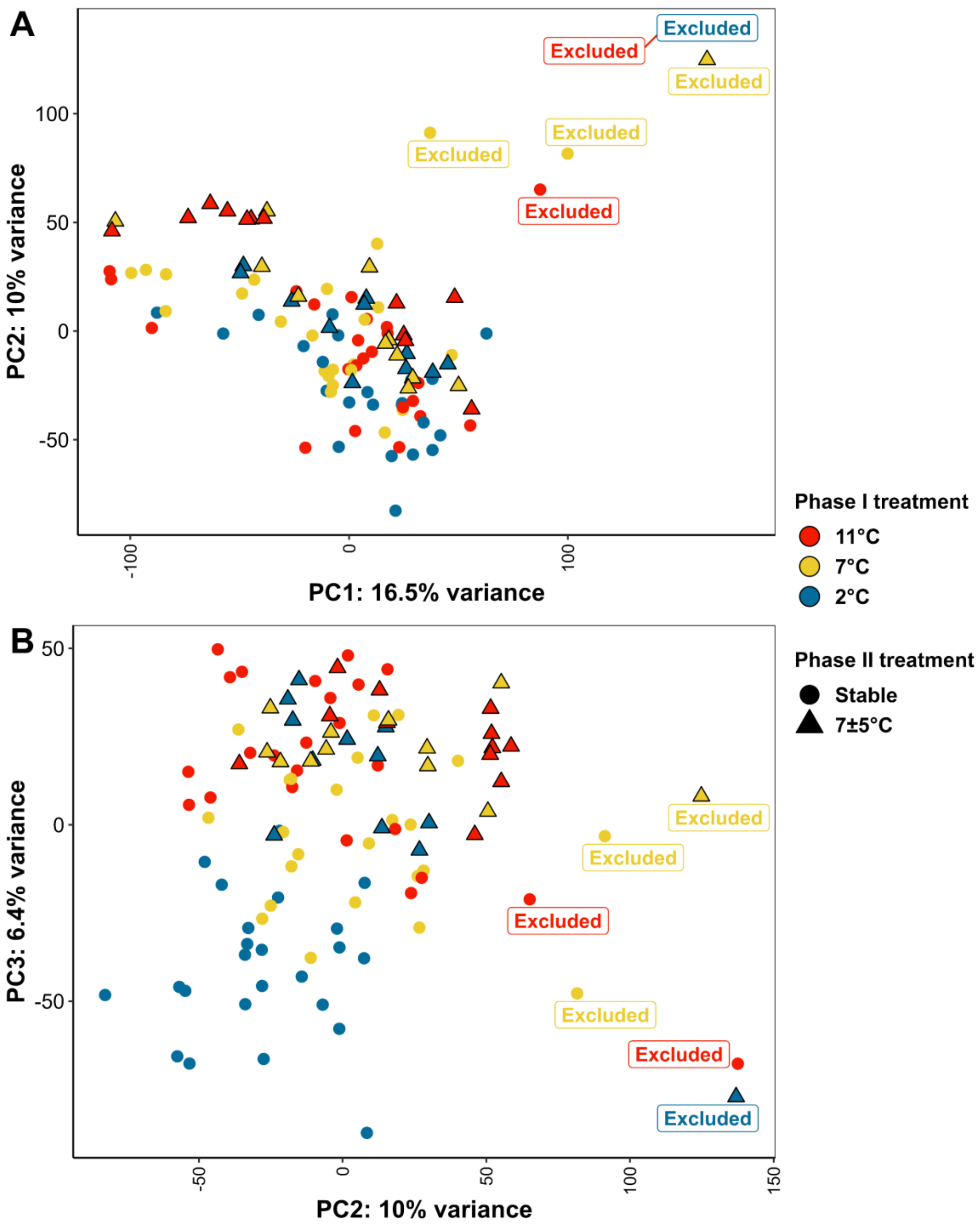


**Figure S3. Principal component analysis (PCA) of RNA-seq samples in Experiment 2 for outlier filtering.** A) principal component one (PC1) and PC2; B) PC2 and PC3. The six samples that derived from the main cluster in all three PCs were excluded.

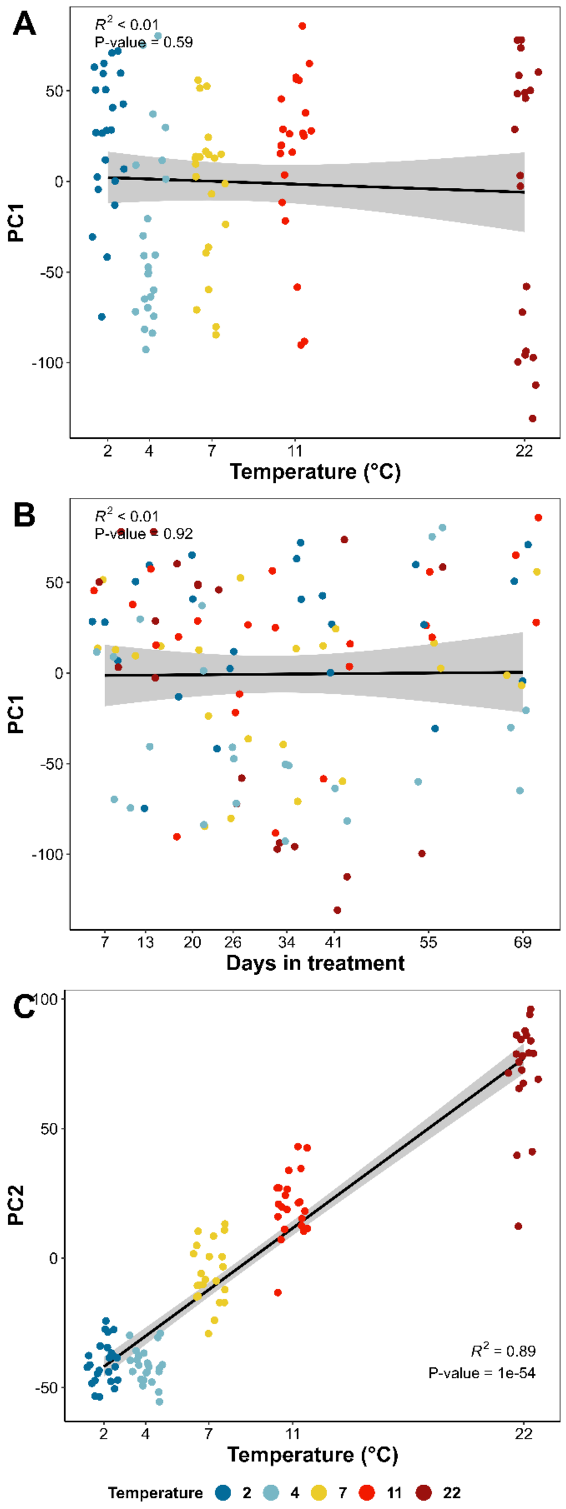


**Figure S4. Correlation analysis of principal component one (PC1) and PC2 with temperature and days in treatment in Experiment 1.** A) Correlation analysis of treatment temperature and PC1; B) Correlation analysis of days in treatment and PC1; C) Correlation analysis of treatment temperature and PC2.

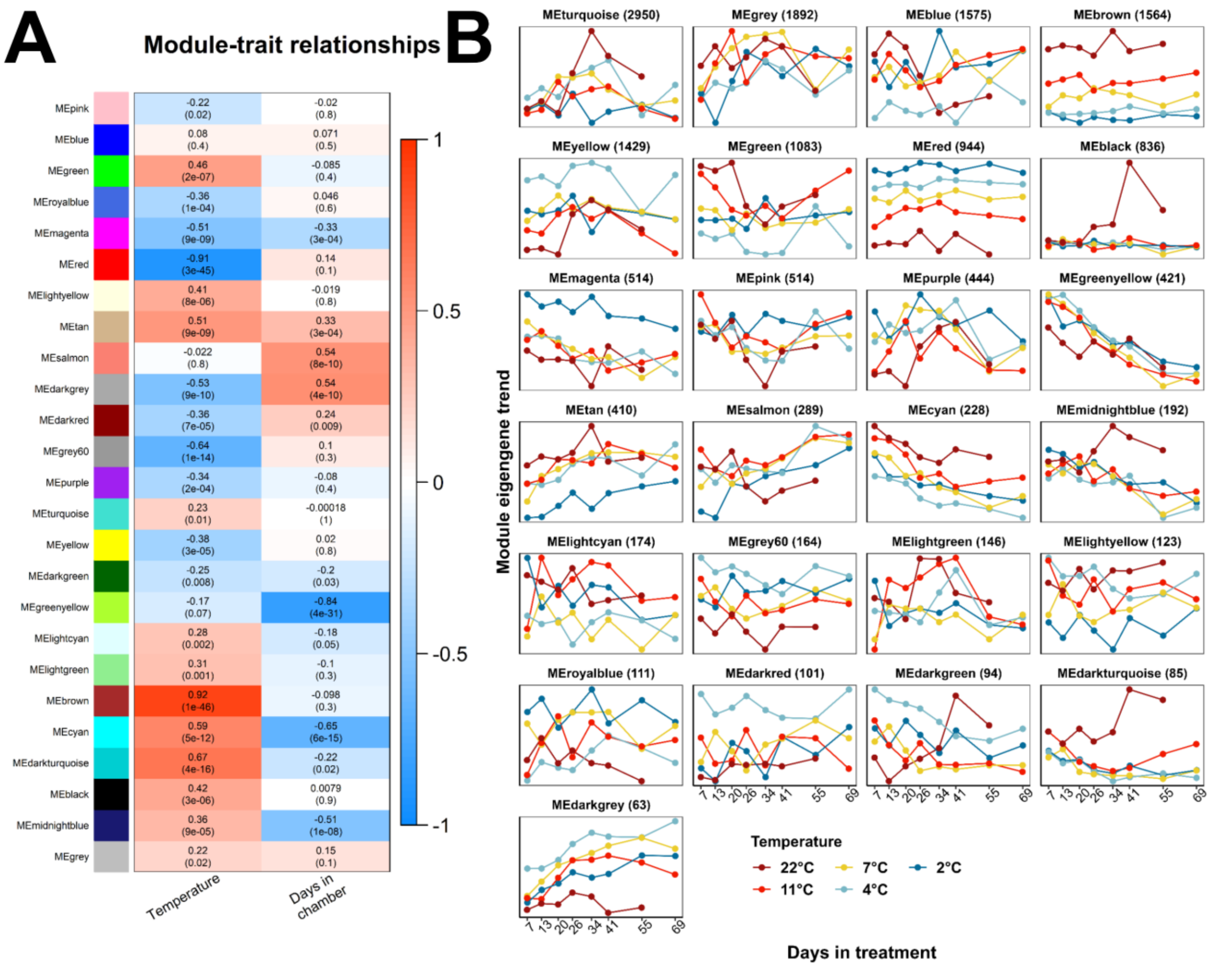


**Figure S5. WGCNA module eigengene correlation analysis and visualization in Experiment 1.** Weighed gene co-expression network (WGCNA) was constructed via 'signed' network type with a soft-thresholding power of 12, and the minimum module size was set as 50. A total of 16,346 genes after low count gene filtering were inputted to WGCNA. The expression of each gene was transformed using variance stabilizing transformation in DESeq2. Module eigengene (ME) is defined as the first principal component (PC1) of the expression of all genes in the module. A) Correlation analysis of ME and treatment temperature and days in treatment; B) Mean ME visualization across the experiment.

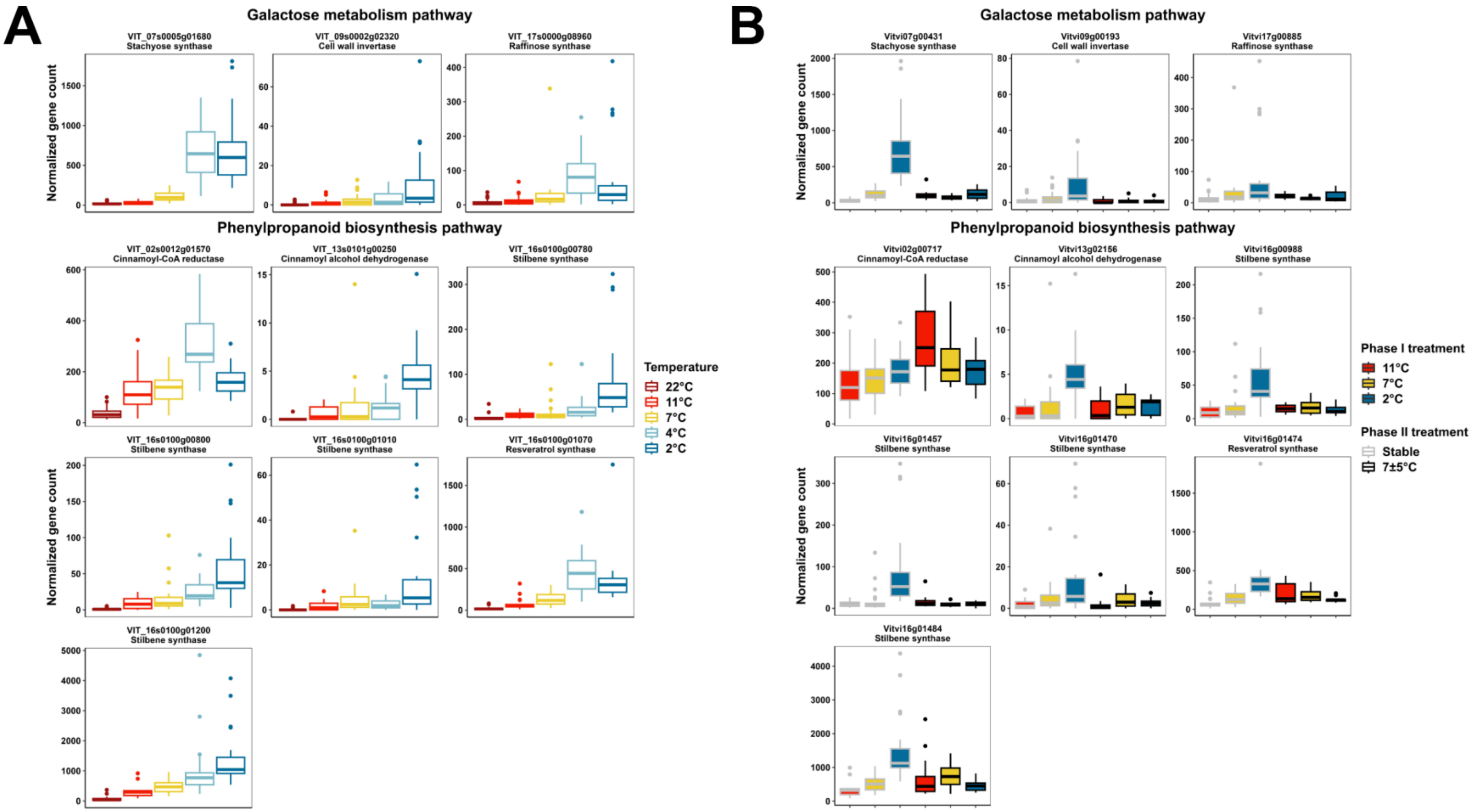


**Figure S6. The expression of top DEGs that negatively correlated with temperature in Experiment 1 in galactose metabolism pathway and phenylpropanoid pathway.** A) Expression in Experiment 1; B) Expression in Experiment 2. The ranking of DEGs is based on LFC in the correlation analysis of gene expression and temperature, and only the top 200 genes were scouted. The expression of genes is normalized using DESeq2.

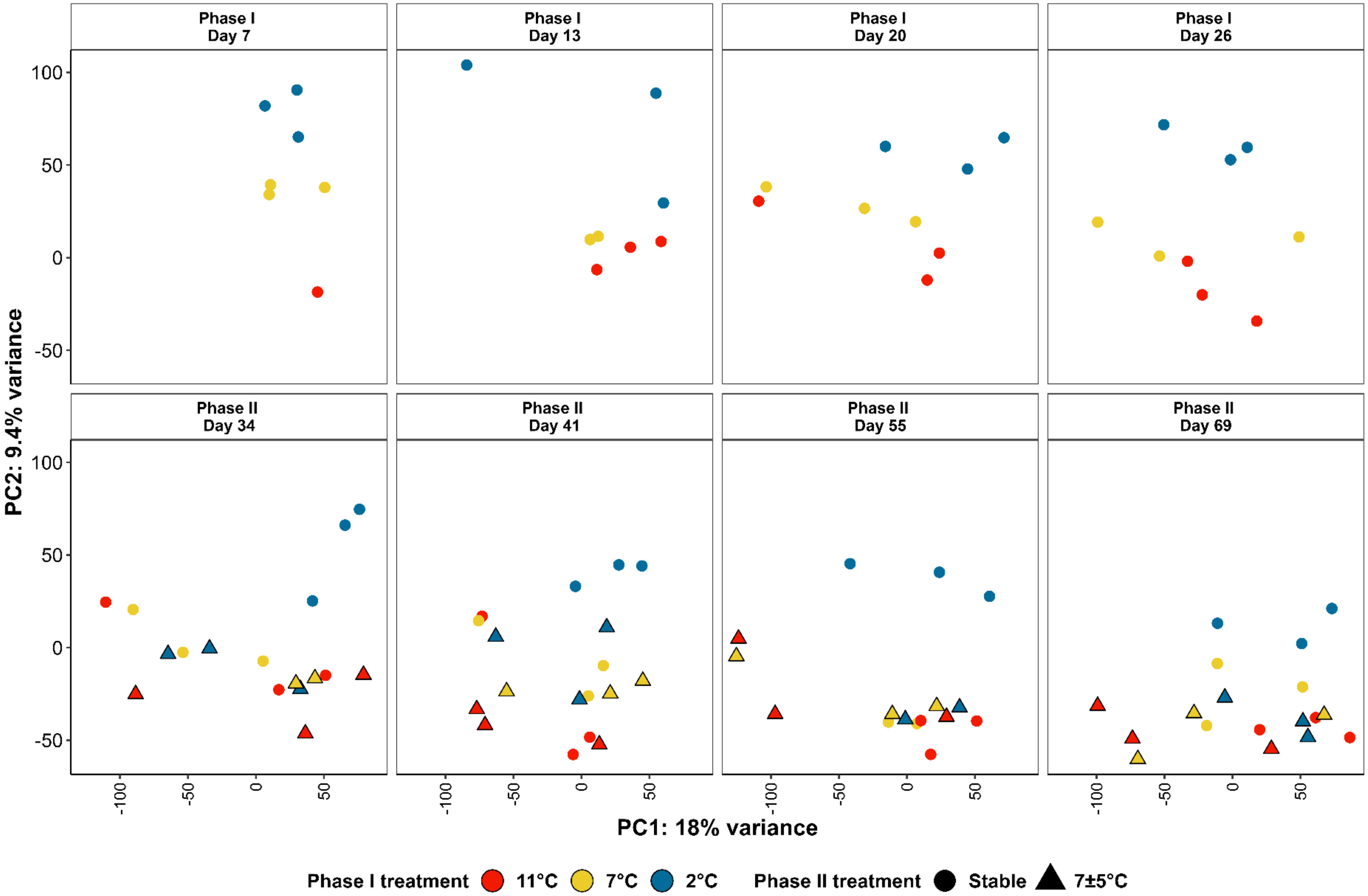


**Figure S7. Principal component analysis (PCA) of all the genes after outlier filtering in Experiment 2.** Only principal component one (PC1) and PC2 are shown.

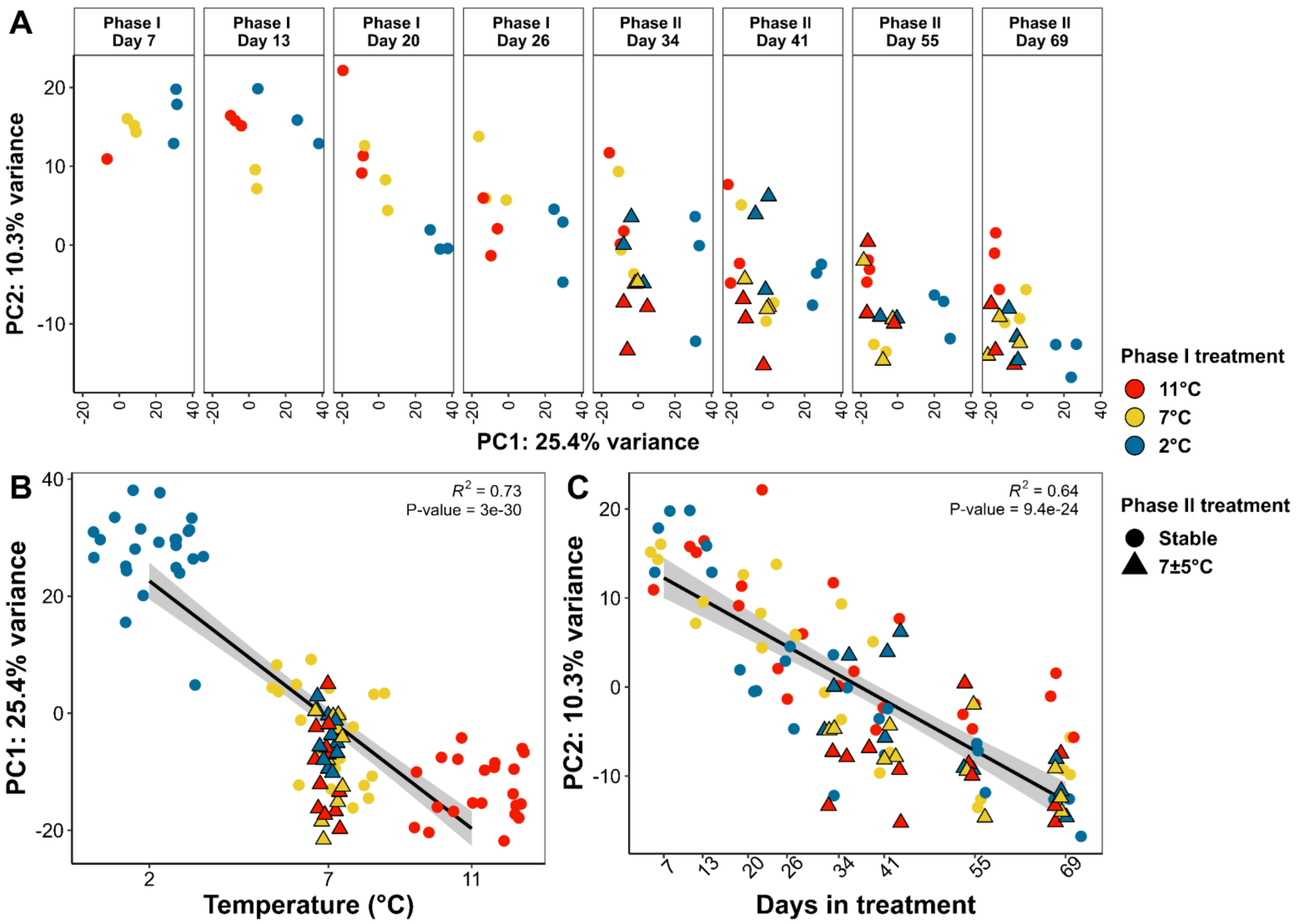


**Figure S8. Principal component analysis (PCA) of the 2,856 temperature-responsive genes identified in Experiment 1 in the samples in Experiment 2.** A) principal component one (PC1) and PC2 at each sample collection; B) Correlation analysis of PC1 and mean treatment temperature; C) Correlation analysis of PC1 with days in treatment.

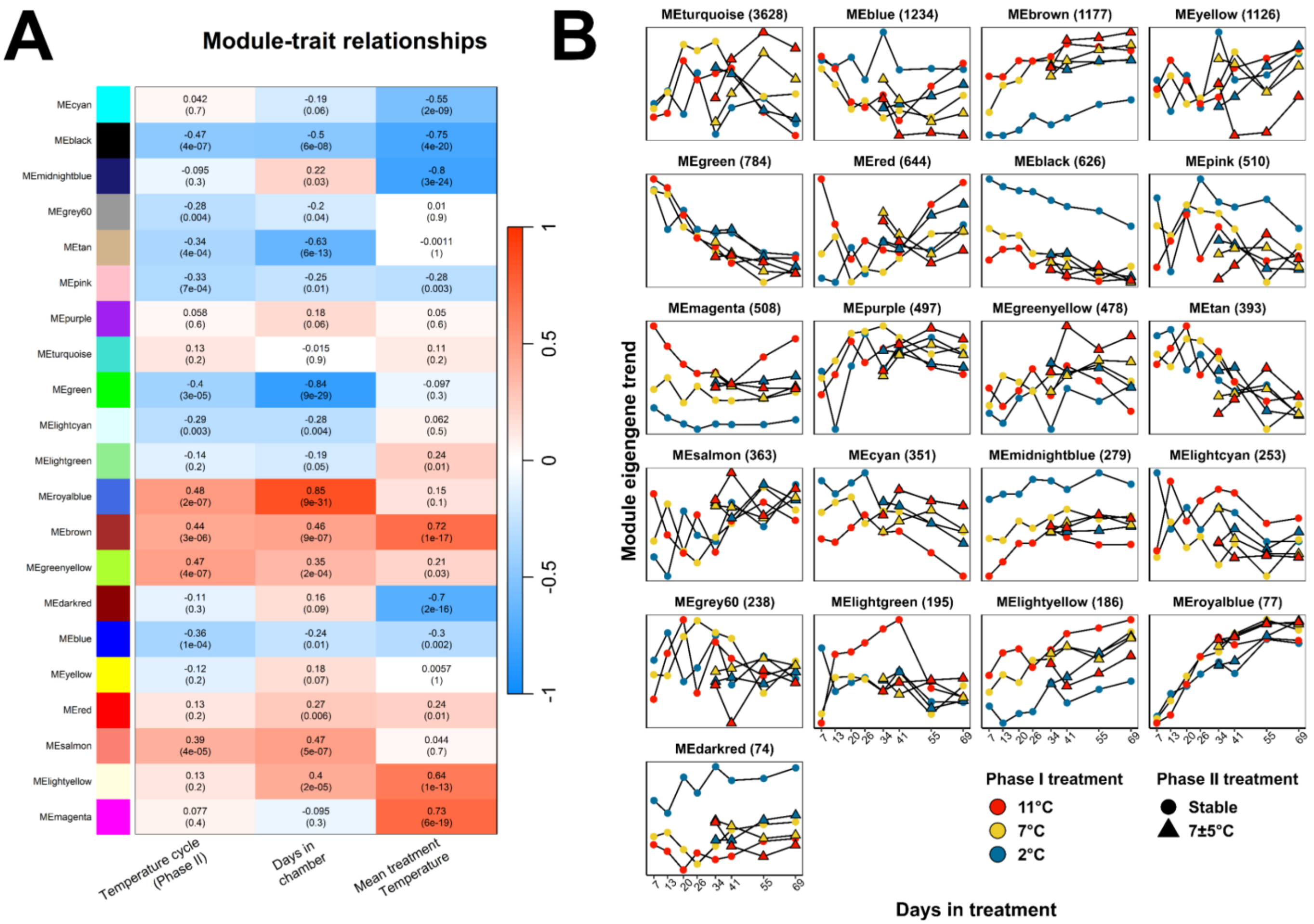


**Figure S9. WGCNA module eigengene correlation analysis and visualization in Experiment 2.** Weighed gene co-expression network (WGCNA) was constructed via 'signed' network type with a soft-thresholding power of 12, and the minimum module size was set as 50. A total of 16,028 genes after low count gene filtering were inputted to WGCNA. The expression of each gene was transformed using variance stabilizing transformation in DESeq2. Module eigengene (ME) is defined as the first principal component (PC1) of the expression of all genes in the module. A) Correlation analysis of ME and treatment temperatures and days in treatment; B) Mean ME visualization across the experiment.

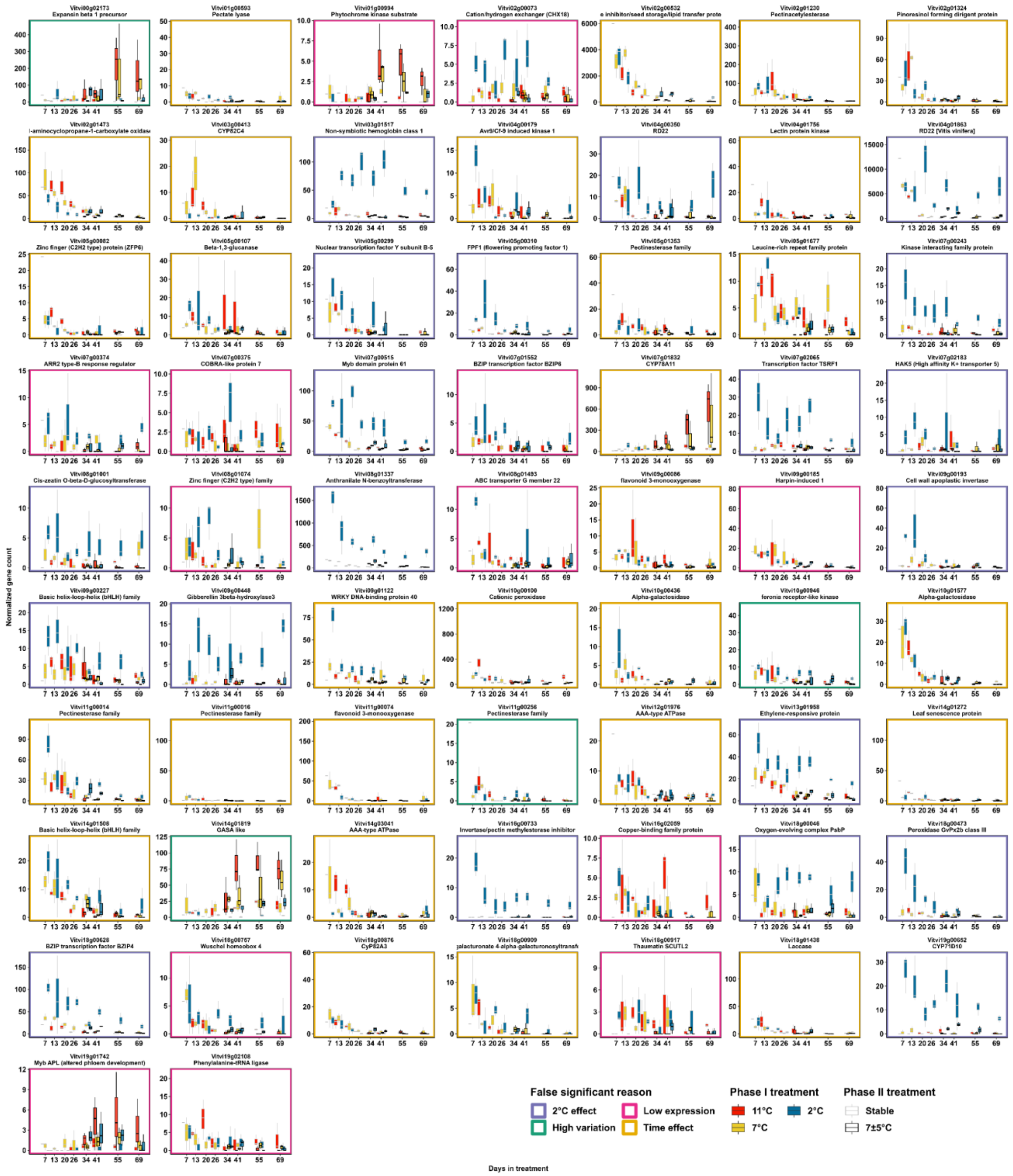


**Figure S10. Differentially expressed genes under temperature cycle treatment in Experiment 2.** Differential expression is determined based on FDR< 0.05 and LFC >2 in the contrast of stable vs. 7±5 °C. The genes that are not functionally annotated in VitisNet are not shown. The expression of genes is normalized using DESeq2.

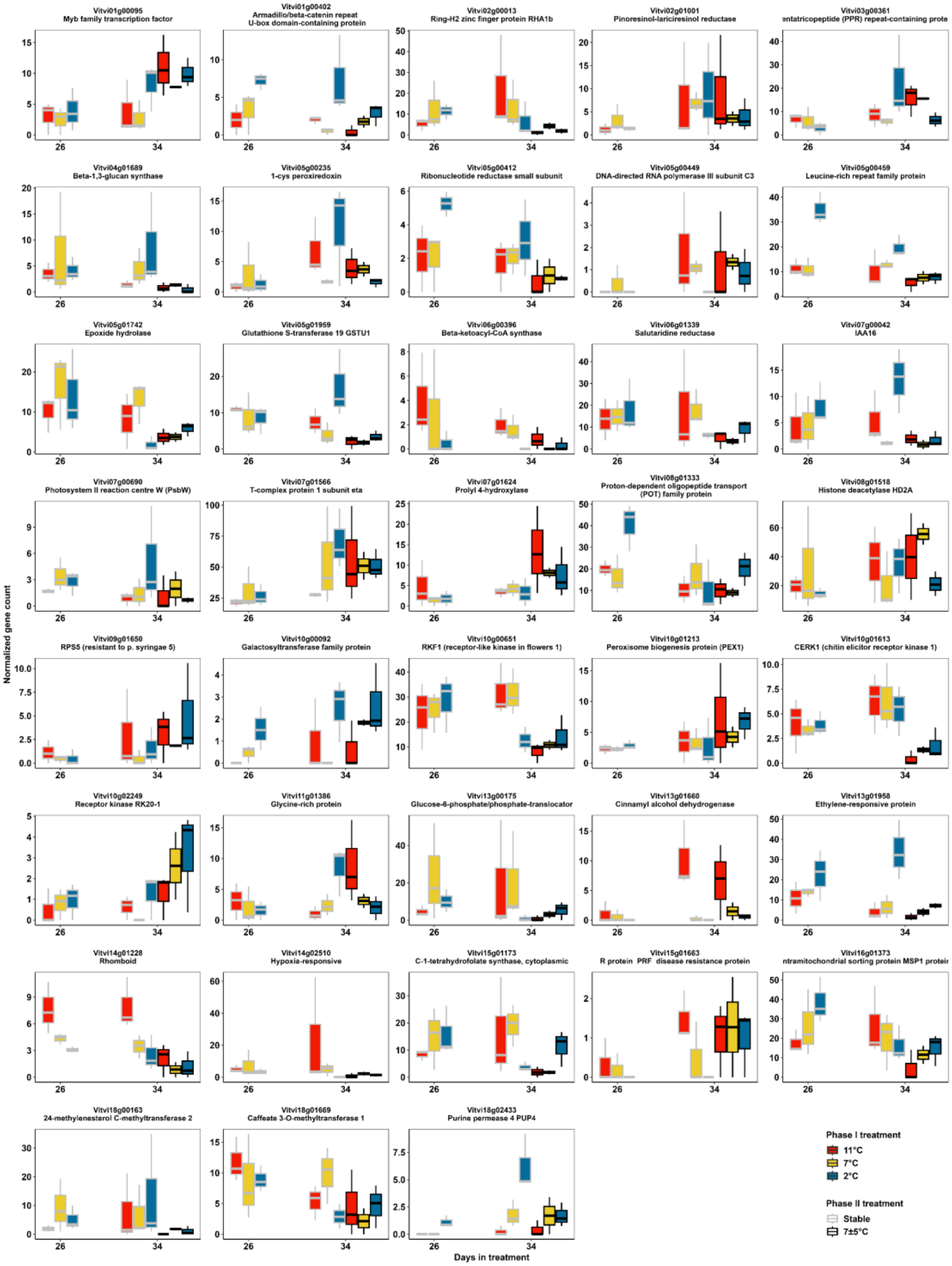


**Figure S11. Differentially expressed genes after eight days of temperature cycling treatment in Experiment 2.** Differentially expressed genes are determined based on shared differential expression at FDR< 0.05 and LFC >1 in the contrasts of phase I 2 °C on 26 d vs. phase I 2 °C × phase II 7±5 °C on 34 d, phase I 7 °C on 26 d vs. phase I 7 °C × phase II 7±5 °C on 34 d and phase I 11 °C on 26 d vs. phase I 11 °C × phase II 7±5 °C on 34 d. The genes that are not functionally annotated in VitisNet are not shown. The expression of genes is normalized using DESeq2.

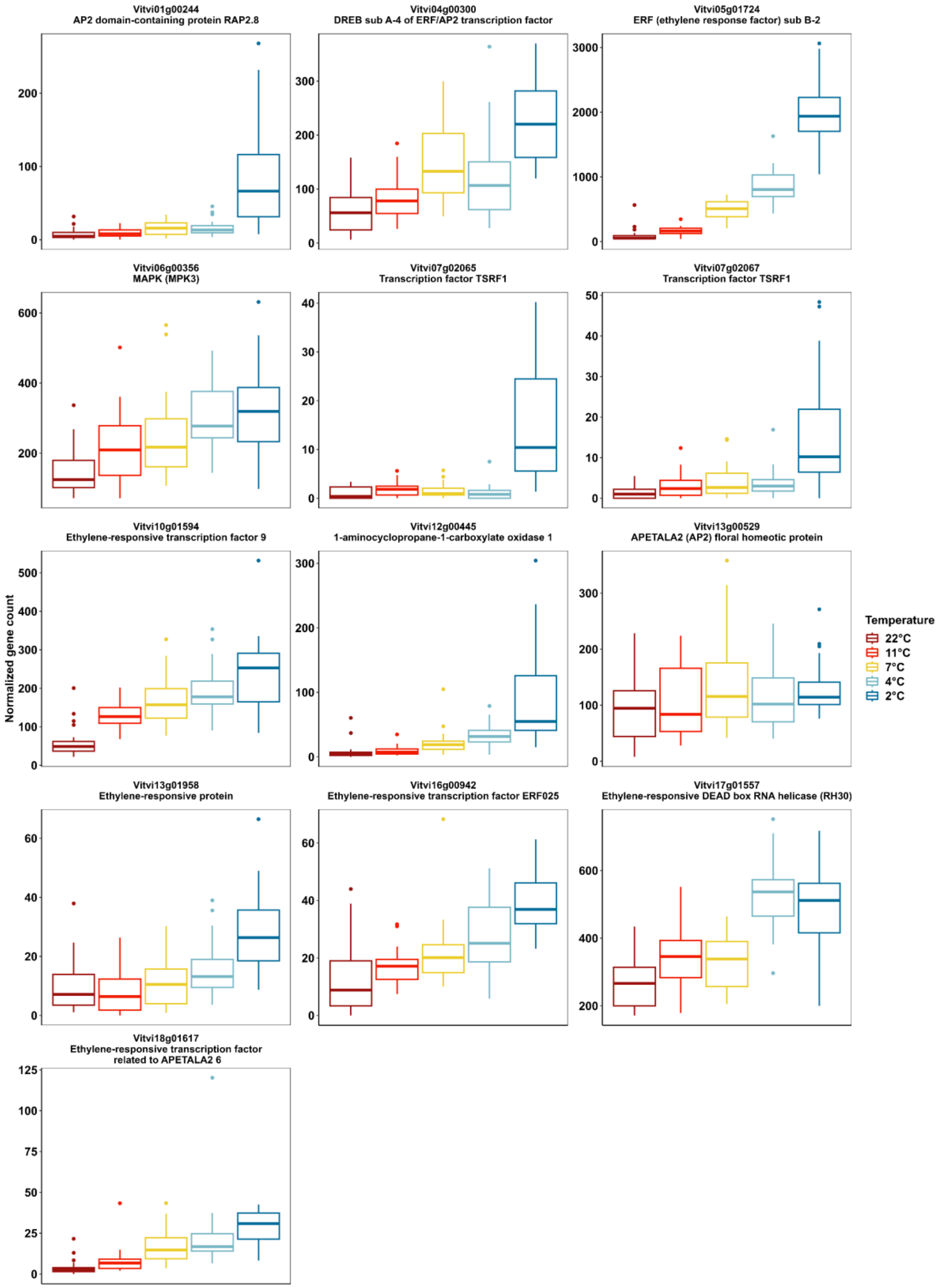


**Figure S12. The expression of all DEGs in ethylene signaling pathway and *ERFs* that negatively correlated with temperature in Experiment 1**. The expression of genes is normalized using DESeq2.

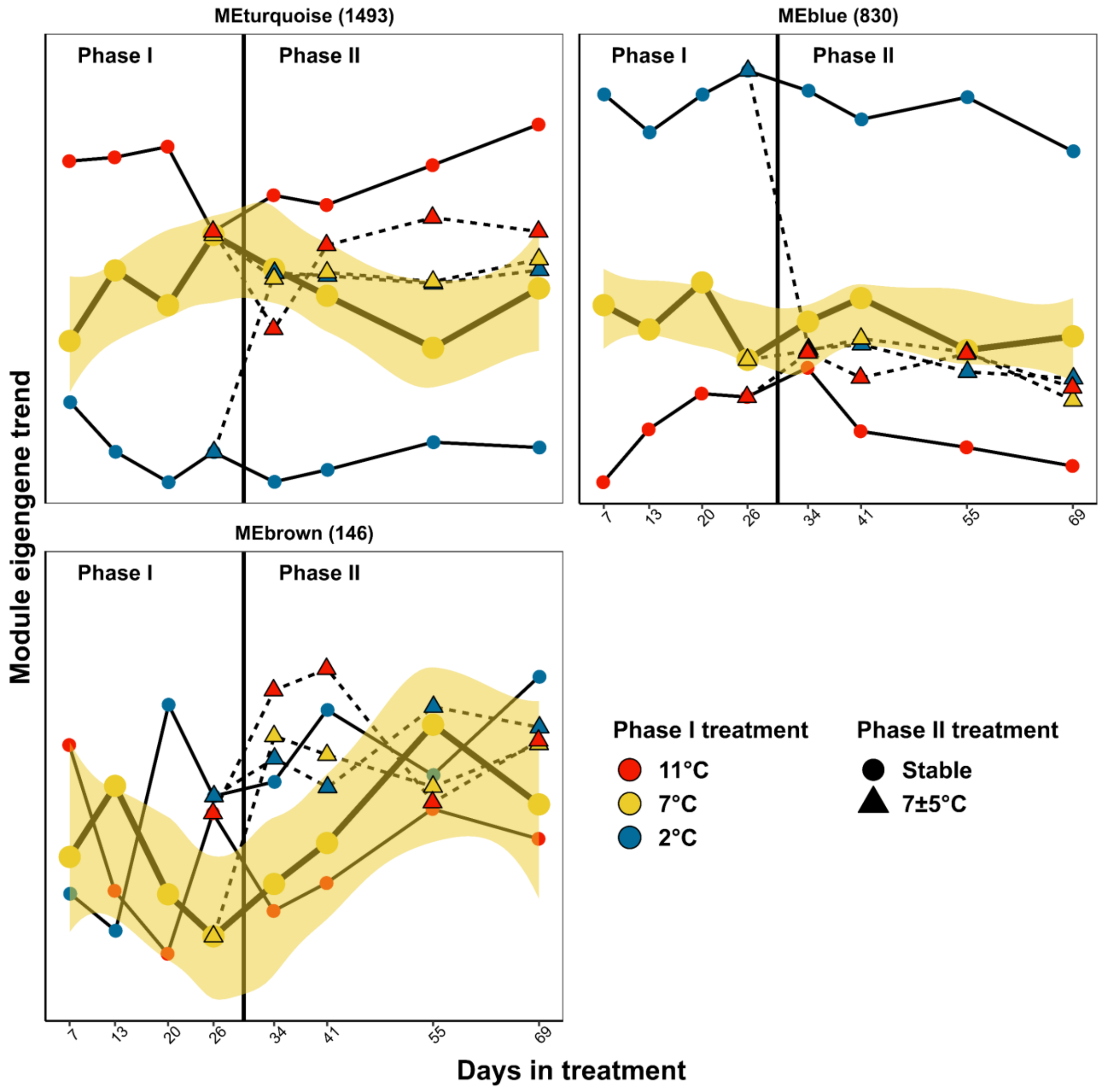


**Figure S13. Module eigengenes in the WGCNA of Experiment 2 sample using only temperature-responsive genes identified in Experiment 1**. Weighed gene co-expression network (WGCNA) was constructed via 'signed' network type with a soft-thresholding power of 12, and the minimum module size was set as 50. A total of 2,856 genes were inputted to WGCNA. The expression of each gene was transformed using variance stabilizing transformation in DESeq2. Module eigengene (ME) is defined as the first principal component (PC1) of the expression of all genes in the module.

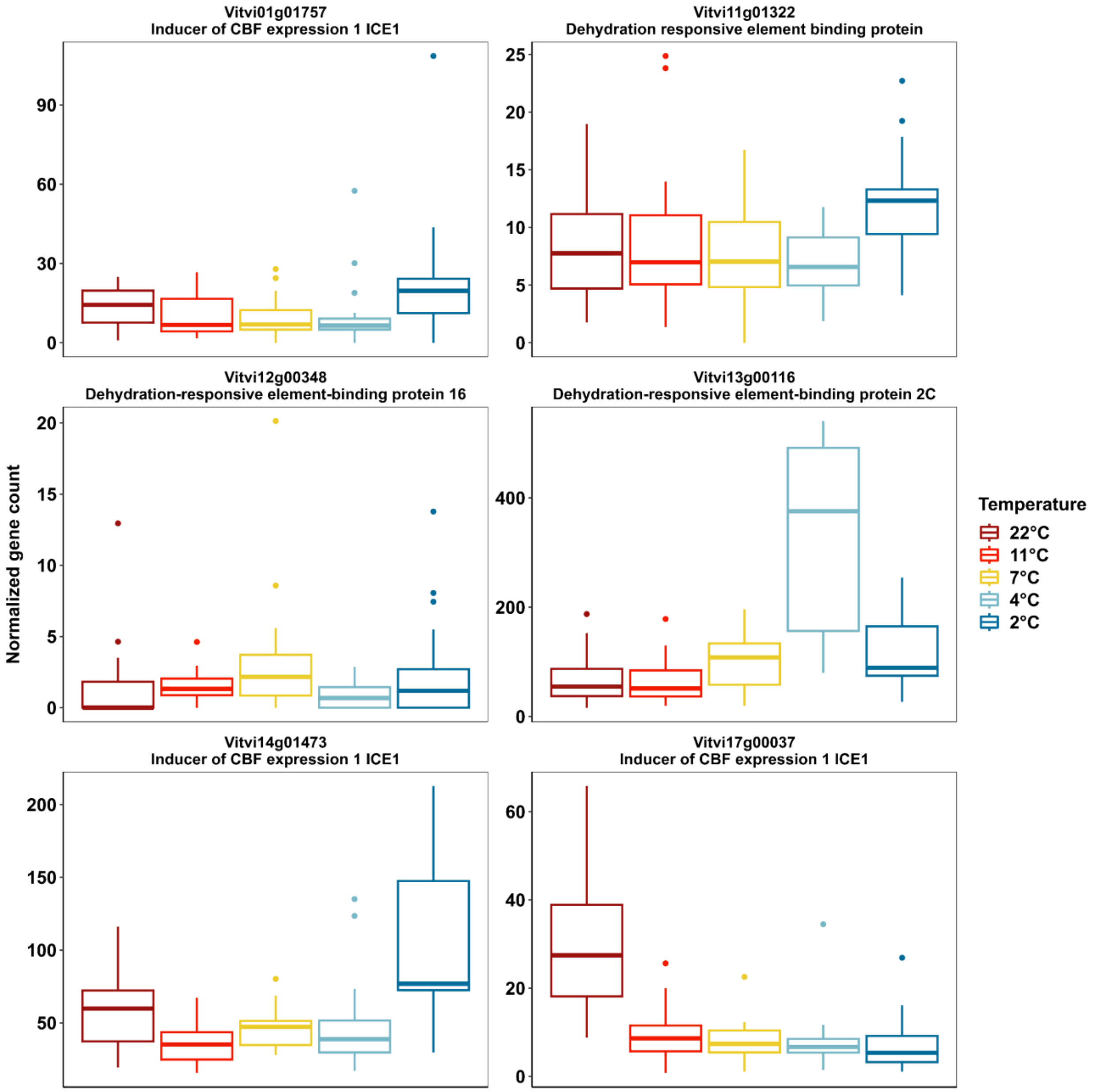


**Figure S14. The expression of ICE and CBF/DREB genes that passing low count filtering in Experiment 1**. The expression of genes is normalized using DESeq2.

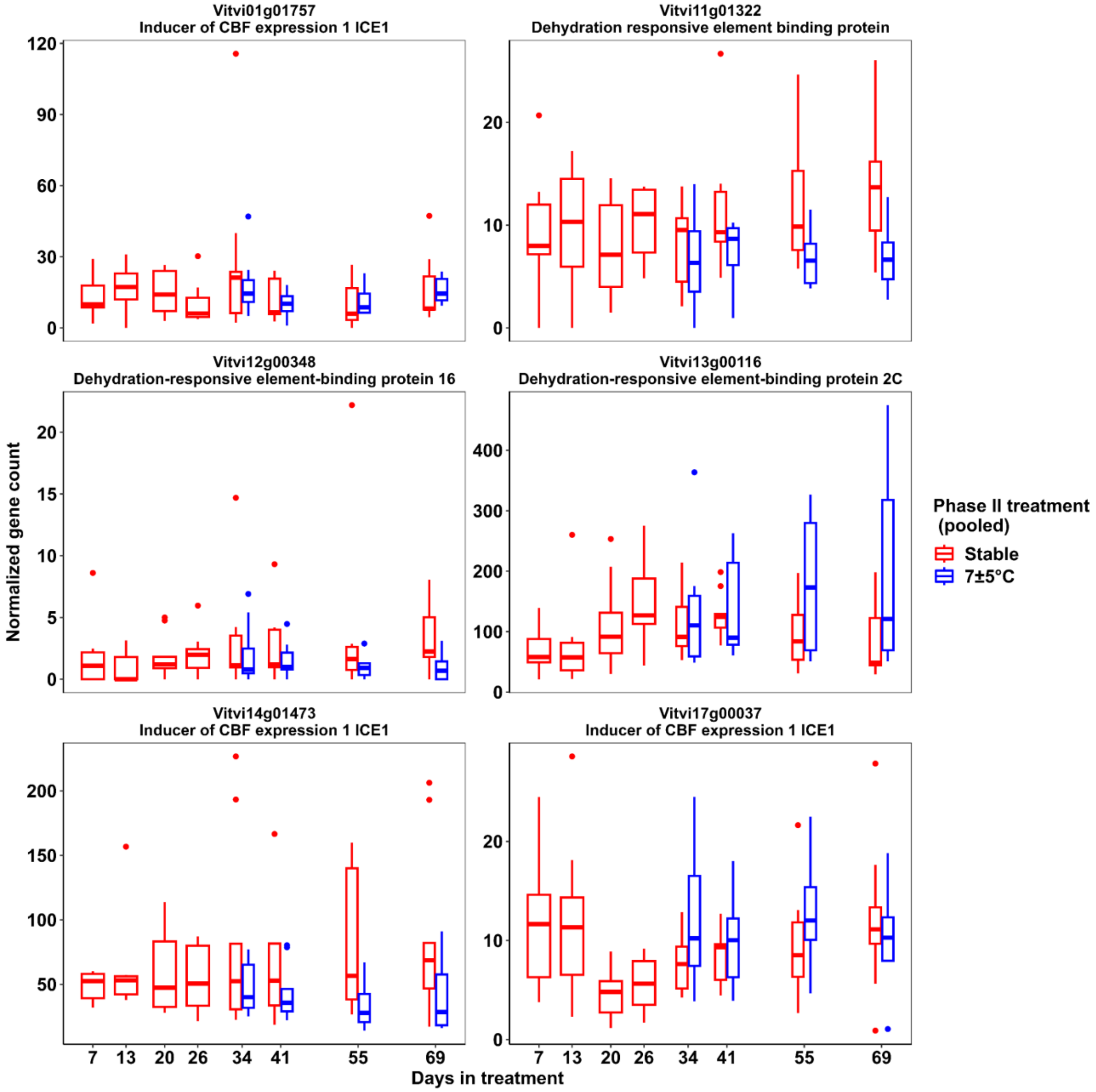


**Figure S15. The expression of ICE, DREB and CBF genes that passing low count filtering in Experiment 2.** The expression of genes is normalized using DESeq2. The gene expression is pooled based on phase II treatment.
